# *Arabidopsis* natural variation induces complex transcriptomic heterochronies at the floral transition and perturbs the coupling between leaf and flower development by an unexpected genetic control

**DOI:** 10.1101/2024.08.12.607587

**Authors:** Dieudonné Sana, Kristianingsih Ruth, Laine Stéphanie, Jesson Béline, Vidal Véronique, Wells Rachel, Morris Richard, Besnard Fabrice

## Abstract

Plant development may be viewed as a sequence of tightly orchestrated events in space and time. The coordination of developmental stages gives rise to various organs, such as leaves or flowers. For instance, in *Arabidopsis thaliana*, leaf/bract development ceases once the first flower emerges. Unravelling the mechanisms of this regulation is, therefore, key for understanding how developmental programs are coordinated during the floral transition. In this study, we take advantage of a previously overlooked natural variation that desynchronizes bract repression from the initiation of the first flowers. We found that prolonged bract formation after the floral transition depends on complex genetic interactions, involving at least four loci. Surprisingly, none of these loci contained the floral identity genes previously implicated in bract repression, suggesting that other genes are involved in the synchronisation of bract and flower programs. Furthermore, although ectopic bracts in inflorescences have been interpreted as a prolonged vegetative state, time-series transcriptomics and curve registration revealed a more complex scenario whereby many genes desynchronize in terms of expression between these developmental programs. As a consequence, transcriptional differences peak at the floral transition when bract development is prolonged, affecting a wide variety of biological processes not necessarily associated with vegetative state. A transient increase in transcriptome divergence has been proposed to account for morphological variation between species in animals and plants under the “inverse hourglass” model. Our results suggest that such a model could also explain the sensitivity of certain developmental transitions to phenotypic variation within species, as reported here for the floral transition.

## Introduction

The development of living organisms generally proceeds through distinct temporal phases. As illustrated by insects developing from embryo to adult through several larval stages and metamorphosis, development is often described as stepwise ontogenetic sequences (Baressi and Gilbert 2023). Some transitions are sharp, requiring the precise coordination of different organs’ developmental programs to generate new structures and ultimately form the organism (Furlow and Neff 2006; Truman 2019; Yamaguchi 2021). At the molecular level, these transitions involve highly dynamic changes in gene expression. Changes in the relative timing of these developmental events, or heterochrony, are recognized as an important drivers of morphological evolution (Alberch et al. 1979; Keyte and Smith 2014). Hence, understanding how developmental timing is controlled from gene expression to organ formation is key to understanding both phenotypic robustness and evolution.

Flowering is a major developmental and morphological transition, transforming plant architecture. In most species, flowers emerge along stems from previous flowerless parts, without any intermediate floral forms. Very often, the start of flower production is also phased with other phenotypic changes. The floral transition thus provides an excellent developmental system to investigate the mechanisms of phenotypic robustness, involving precise spatio-temporal coordination events.

In several species, flower production coincides with the cessation of leaf production, resulting in flower-bearing stems devoid of leaves. In botany, a leaf associated with a single flower is called a bract (Penin 2008; Prenner et al. 2009). A bractless (i.e. leafless) inflorescence is a typical trait of the *Brassicaceae* family, which includes the model plant *Arabidopsis thaliana* (Penin 2008). The presence of bracts in angiosperm inflorescences varies widely among species, showing that the coordination of the leaf program with reproductive development has been independently and recurrently modified during evolution (Mach 2010).

In this study, we use the floral transition in *A. thaliana* as a model system to study how two developmental programs, leaf and flower, are coordinated in a short time window. For instance, several studies have explored how leaf development is halted precisely when the first flower is formed in this species.

First, morphological deformations in the inflorescence meristems (Kwiatkowska 2006) and localised gene repression (e.g., SHOOT MERISTEMLESS [STM]) or activation (e.g., *AINTEGUMENTA [ANT]* or *FILAMENTOUS FLOWER [FIL]*) indicate that a bract domain forms before each flower but fails to outgrow (Long and Barton 2000; Dinneny et al. 2004). This domain is later incorporated in floral tissues, remaining cryptic (Heisler et al. 2005; Goldshmidt et al. 2008). Hence, the absence of a bract is due to its early repression when flower development is initiated.

Second, several genes have been implicated in bract repression through identification of mutants that restore bract formation. These genes, mostly transcriptional regulators, are all functionally associated with the floral meristem identity (FMI) pathway (Siriwardana and Lamb 2012), including the central flower regulator *LEAFY* (*LFY*) (Schultz and Haughn 1991; Weigel et al. 1992). The FMI pathway specifies floral rather than shoot identity (Weigel 1995) and its impairment leads to flower-to-shoot conversions by restoring vegetative characteristics into flowers (altered floral organ identity and phyllotaxis, internode elongation, indeterminacy), including bract outgrowth (see **Table S1**). Furthermore, the genetic ablation of flowers induces bract outgrowth (Nilsson et al. 1998). This suggests that many genes within the FMI regulatory network redundantly act as bract repressors. In all these studies, bract de-repression is consistently accompanied by pleiotropic phenotypes affecting flowers, branching or flowering time.

Conversely, only few genes have been identified as promoting bract outgrowth, such as *JAGGED* (Dinneny et al. 2004; Ohno et al. 2004), *AINTEGUMENTA* (*ANT*) and *AINTEGUMENTA-LIKE6* (*AIL6*) (Manuela and Xu 2024 Mar 28) and their misexpression also produce pleiotropic phenotypes in addition to bract development. Bracts can also be induced in wild-type *A. thaliana* under light treatments enriched in far red light, together with flower-shoot chimeras (Hempel and Feldman 1995; Hempel et al. 1998). The authors proposed a “conversion” model, where a young branch meristem (already subtended by a leaf) can be converted (here upon photo-induction) into a flower that inherits this pre-formed leaf, interpreted then as a bract. However, the underlying genetic mechanisms also remain unclear.

In summary, the current literature suggests a straightforward explanation for the precise coupling of bract repression with formation of the first flower in *A. thaliana:* this arises from the bract-repressing activity of the FMI pathway acting in the flower. The molecular mechanisms involved from flower to bract remain unclear but they are likely indirect and/or non-cell autonomous, as FMI genes are mostly expressed in the developing flower and not in the bract domain. Since bracts are almost identical to leaves produced during the preceding vegetative phase (Dinneny et al. 2004), de-repressed bracts have sometimes been interpreted as the persistence of a vegetative program (Hempel et al. 1998). Bract de-repression and improper establishment of flower identity are therefore considered as two sides of the same coin.

In this study, we provide new data showing that, in natural populations, bract repression and flower development can be uncoupled. We report the transient presence of bracts with wild-type flowers precisely at the floral transition in wild *A. thaliana* accessions. By combining precise phenotype characterization, quantitative genetics, genomics, and time-series transcriptomics specifically in meristem tissues, we investigated this change in the timing of flower and leaf formation at developmental, genetic and transcriptomic levels. Altogether, our results provide two new main contributions to our understanding of the mechanisms coordinating leaf and flower development. First, bract development can be prolonged without affecting the main FMI pathway genes. Second, this prolonged bract formation at the floral transition does not correspond to a global “juvenile” shift in the transcriptome and correlates with more complex transcriptomic heterochronies. In line with the ‘inverse hourglass model,’ we discuss how transcriptional divergence, resulting from the ongoing adaptation of flowering timing, may lead to phenotypic variation at different evolutionary scales.

## Results

### Bract repression is not strictly synchronised with the floral transition in the natural accession *Tsu-0*

Like in *A. thaliana* and its reference accession (*Col-0*) (**Fig. 1A**), the typical ebracteate flower (i.e. without bracts) of *Brassicaceae* generally appears as soon as the first flower is produced, indicating synchronization of bract repression with the onset of the flower program. However, in a number of *Brassicaceae* species, bract repression is delayed for several rounds of flower formation, generating bracteate flowers (i.e. with bracts) at the base of their flowering branch (a raceme) (Endress 2006; German et al. 2023): some species display bracts up to the first half of the raceme (e.g. **Fig. S1A,C**), while others only have one or two bracts at the base (**Fig. S1B,D** or e.g. in Al-Shehbaz 2015).

**Figure 1:**
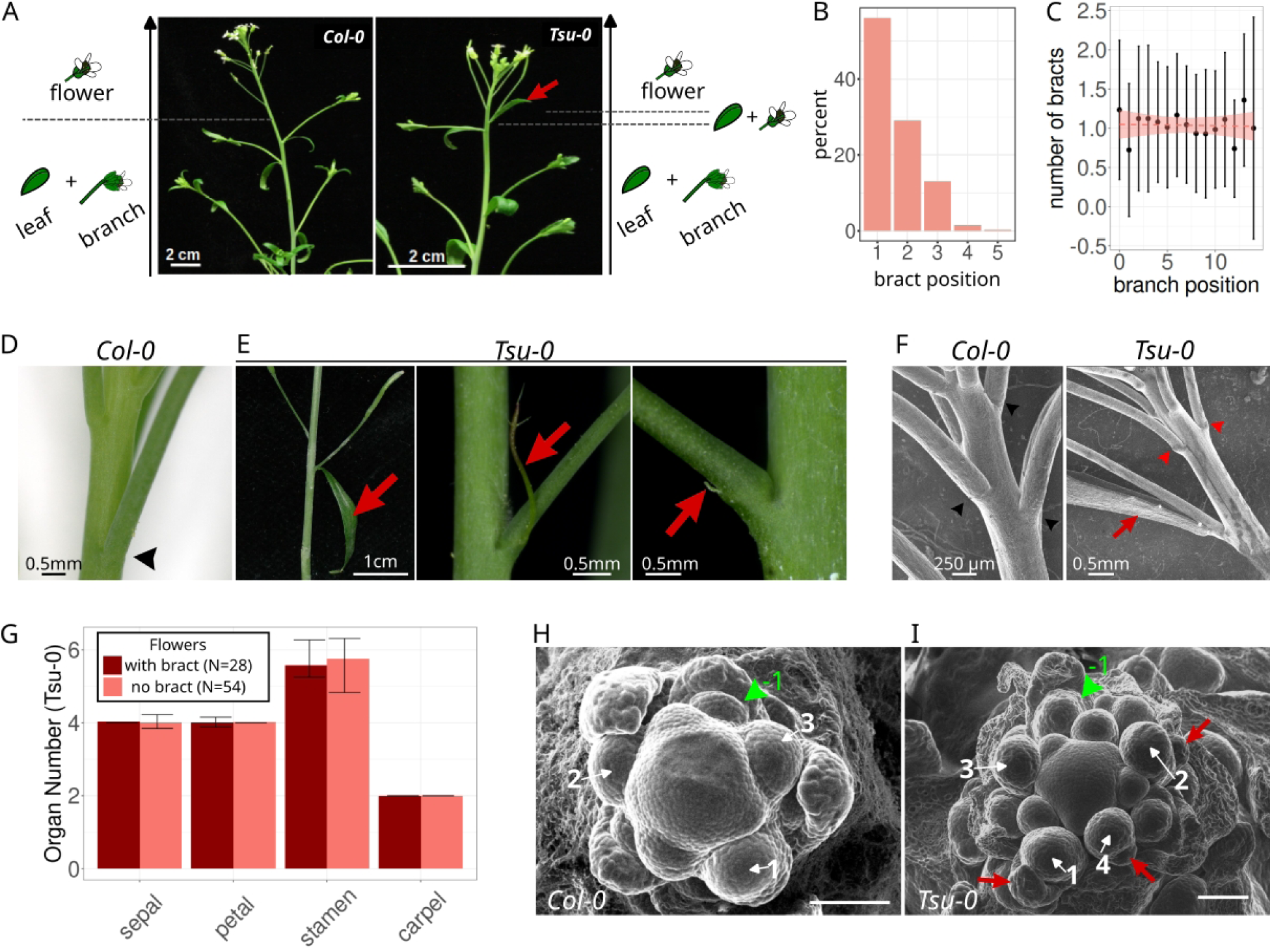
Desynchronization of leaf repression from flower formation in the natural *Tsu-0* accession of *Arabidopsis thaliana*. **A,** lateral views of the main stem of *Col-0* (left) and *Tsu-0* (right) accessions showing the transition from branches (at the bottom) to flowers (to the top). The red arrow points to a bract (a leaf) subtending the first flower in *Tsu-0*. Cartoons on the side summarize the structures produced at each node. **B**, Distribution (as percent) of the position of bracts in *Tsu-0* (N plants = 50, n bracts = 337) in the main stem and cauline branches, where n°1 is the first flower after floral transition. **C,** Number of bracts on the different branches of a *Tsu-0* inflorescence (dot is the mean, vertical bars are sd). Branch positions are numbered with increasing numbers from the base to the top of the main stem, which is noted as 0 by convention (N plants = 90). A linear regression (horizontal red dotted line with standard error shaded in red) indicates a stable average value of 1 bract on each branch. **D,** *Col-0* floral pedicels do not show any sign of bract development (black arrowhead). **E**, Variable bract development (red arrows) observable in *Tsu-0*, from true leaves (left) to filamentous or small rudimentary structures at the base of the floral pedicel. **F**, Scanning electron microscopy of floral pedicels at the floral transition in *Col-0* (left) and *Tsu-0* (right). In *Col-0*, the pedicel is a regular, naked cylinder at the base (black arrow) while in *Tsu-0*, there is a bract on the first flower (red arrow) and swollen base of pedicels on the two following flowers (red arrowhead). **G**, Number of floral organs in flowers with (dark red) or without (light red) bract in *Tsu-0* plants. (**H,I**) Scanning electron microscopy of the main meristem of plants captured right at the transition to flowering in *Col-0* (**F**) and *Tsu-0* (**G**). The first flowers after the transition are pointed by white arrows and ordered with increasing numbers in their order of production inferred from size and phyllotactic positions. Green arrowheads: cauline branch (axillary meristem and a cauline leaf), −1 indicates the last branch before floral transition. Red arrows: bracts. In *Col-0,* no bract is visible on the adaxial side of the developing flowers.

Likewise, we discovered that the natural *Tsu-0* accession in *A. thaliana* displays bracts on the first one to five flowers of the raceme (**Fig. 1A,B**), in the main stem and cauline branches at the same frequency (**Fig. 1C**). A gradient of bract outgrowth can be observed in *Tsu-0*, from fully developed bracts resembling the cauline leaves associated to branches, up to filamentous, short rudimentary structures (**Fig. 1D,E**). Scanning electron microscopy (SEM) showed that a mild swelling of the peduncle basis could sometimes occur in *Tsu-0*, unlike in *Col-0* (**Fig. 1F**). Both bracteate and ebracteate *Tsu-0* flowers were normal, containing the invariant organ number typical of each whorls. Thus, developmental variation is restricted to the bract (**Fig. 1G**). To exclude that these leaves could be secondary outgrowth from mature floral peduncles, we imaged the emergence of the first flower by SEM. In *Col-0*, the last lateral meristem subtended by a cauline leaf was immediately followed by ebracteate flowers (**Fig. 1H**). In *Tsu-0*, the first flowers produced after the last branch displayed bracts or rudimentary bracts (**Fig. 1I**), recapitulating the gradient of bract forms observed in mature stems. Their abaxial position and their precocious development also suggest that these structures developed from the cryptic bract, that developed despite the onset of flower formation.

Since *Tsu-0* is a wild accession, we wondered whether a ‘conversion’ model as proposed by Hempel and Feldman (1995; 1998) for other wild accessions could explain this phenotype. However, phenotyping hundreds of plants did not identify any shoot-flowers chimaeras (**Fig. 1D,F,H**). Also, *Tsu-0* plants did not exhibit the negative correlation between the numbers of bracts and cauline branches (**Fig. 2A**) that would be expected under the “conversion” model. This suggests that *Tsu-0* bracts are not produced by shoot-to-flower conversions.

**Figure 2:**
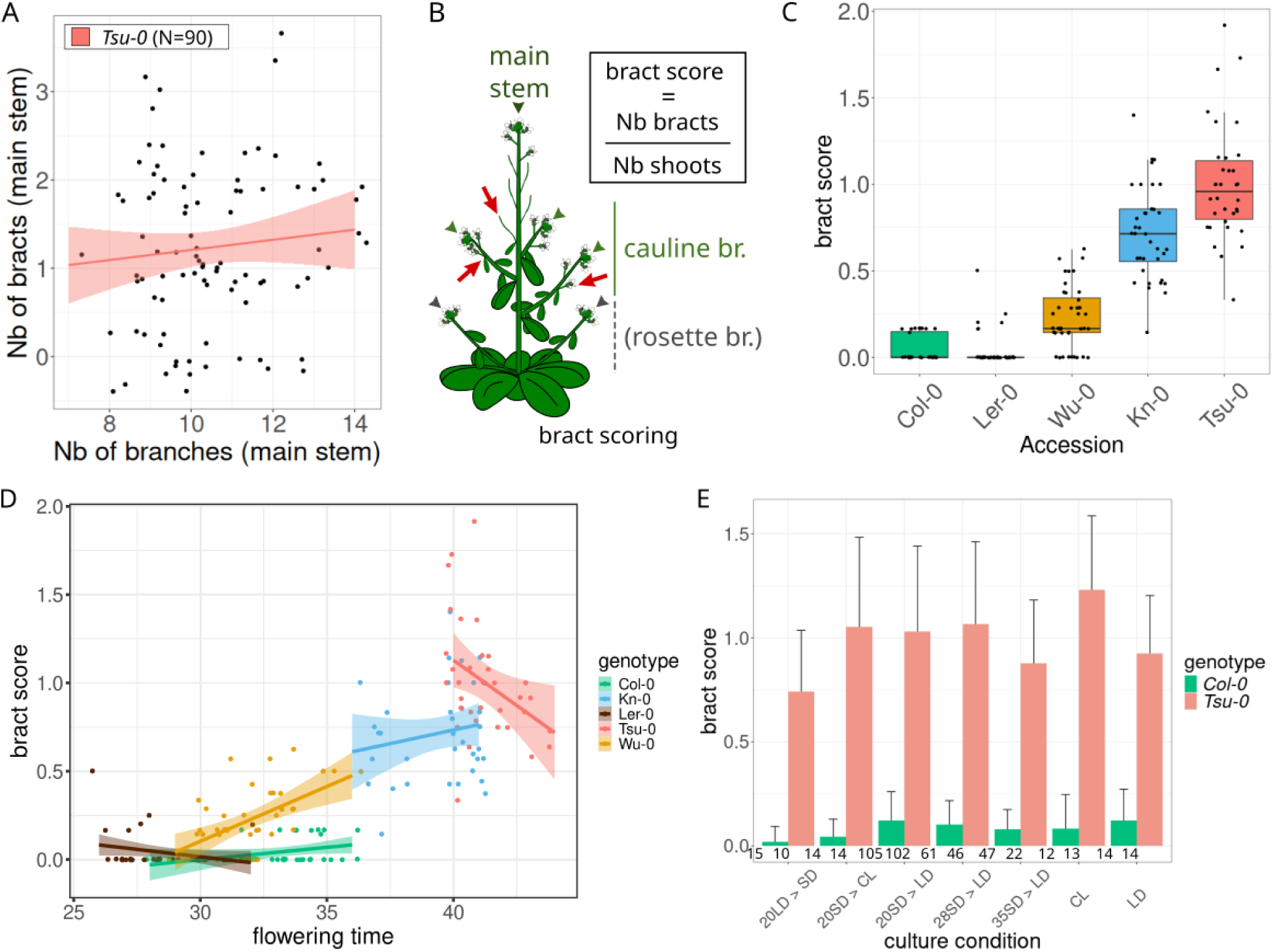
Desynchronization of leaf repression from flower formation in *Tsu-0* and other natural accessions exhibits complex relationships with flowering time and photo-induction conditions. **A**, Correlation study between the number of lateral (cauline) branches and the number of bracts on the main stem in *Tsu-0* (Pearson test, p-value > 0.1), which do not show a negative correlation expected by the “conversion” model (see main text). **B**, Definition of a quantitative bract score for a single plant. **C**, Quantification of basal bracts in different accessions using the bract score. **D**, Correlation study between the bract score and the flowering time (assayed when the first flower opens) in different accessions, labelled with different colours. Each dot is a plant and a linear regression with standard deviation (color shade) is computed for each accession. **E**, Effect of different light regimes on bract scores in *Col-0* and *Tsu-0* plants (green and red, respectively). LD and SD stand for long- and short-day conditions, respectively and CL stands for continuous light (see methods). Numbers (e.g. 20SD > LD) indicate the number of days in the first condition before transfer to the second. The number of plants scored is indicated below each bar.

To summarize, in the natural accession *Tsu-0*, bract repression is no longer synchronized with the onset of flower development. Instead, it is transiently altered to varying and generally decreasing degrees in the first flowers of a branch, resulting in what we have termed ‘basal bracts’.

### Desynchronisation of bract repression with the floral transition is common among Arabidopsis natural accessions

To measure the ability of different genotypes to generate basal bracts, we defined a bract score (**Fig. 2B**) that takes into account several branches (hence several floral transitions) per plant, normalizing by the number of scored branches. Similar to *Tsu-0*, other genetically diverse natural accessions produce basal bracts, albeit at highly variable rates (**Fig. 2C**). We could not identify any correlation between basal bracts and geographic or genetic origins (**Fig. S2**). The bract score shows a high level of intra-genotype variability. Despite this variability, there are clear differences between the basal bracts of different genotypes (**Fig. 2C**). The production of basal bracts is thus a common phenotypic trait among *A. thaliana* natural accessions from different geographical origins and varying genetic backgrounds.

### Basal bract frequency shows complex correlations with flowering time but is not affected by variation in photo-induction

We then explored whether variation in the frequency of basal bracts in natural accessions could be linked to variation in flowering time. Indeed, flowering time is known to be highly variable among natural accessions (Salomé et al. 2011) and depends on both genetic and environmental factors (Maple et al. 2024). We measured both bract score and flowering time in a panel of five accessions and observed a global positive correlation across genotypes (**Fig. 2D**). For instance, *Tsu-0* flowers later than *Col-0* in either absolute time (ie., the number of days from germination) or developmental time (measured by the number of leaves produced), consistently across culture conditions (**Fig. S3**). This suggests that a longer flowering time could promote the alteration of synchrony between bract repression and the onset of flower development. However, within an accession, this correlation can be negative (e.g. in *Ler* and *Tsu-0*), suggesting that the relationships between basal bracts and flowering time could be more complex depending on the genetic backgrounds.

To explore further the effects of varying flowering time in a genetic background, *Col-0* and *Tsu-0* plants were cultured in different photo-induction regimes that vary the flowering time (see methods, see **Fig. S3**). The bract score remains remarkably constant across all tested conditions for both accessions (**Fig. 2E**). This indicates that basal bract formation does not depend either on photoperiod or on the absolute flowering time and rather suggests the importance of intrinsic genetic determinism specific to each accession.

### Delayed bract repression is controlled by multiple QTLs that do not contain genes previously associated to bract development

To analyze the genetics of basal bract formation, we conducted quantitative genetics using crosses between *Tsu-0* and *Col-0* since these two accessions ranked respectively as high and low bract producers (**Fig. 2C**). F_1_ progeny showed intermediate bract scores while F_2_ bract scores spread within the parental range, with a distribution that does not indicate a simple monogenic trait (**Fig. 3A and Fig. S4A**). Bulk Segregant Analysis (BSA) with mapping-by-sequencing identified four quantitative trait loci (QTL): two on chromosome 1 and two on chromosome 5 (**Fig. S4B,C**). The large spread of F_1_ phenotypic values constrained F_2_ plant selection with few plants in the high-score bulk and possibly spurious homozygous in the low-score bulk. These two limits may explain the large intervals obtained.

**Figure 3:**
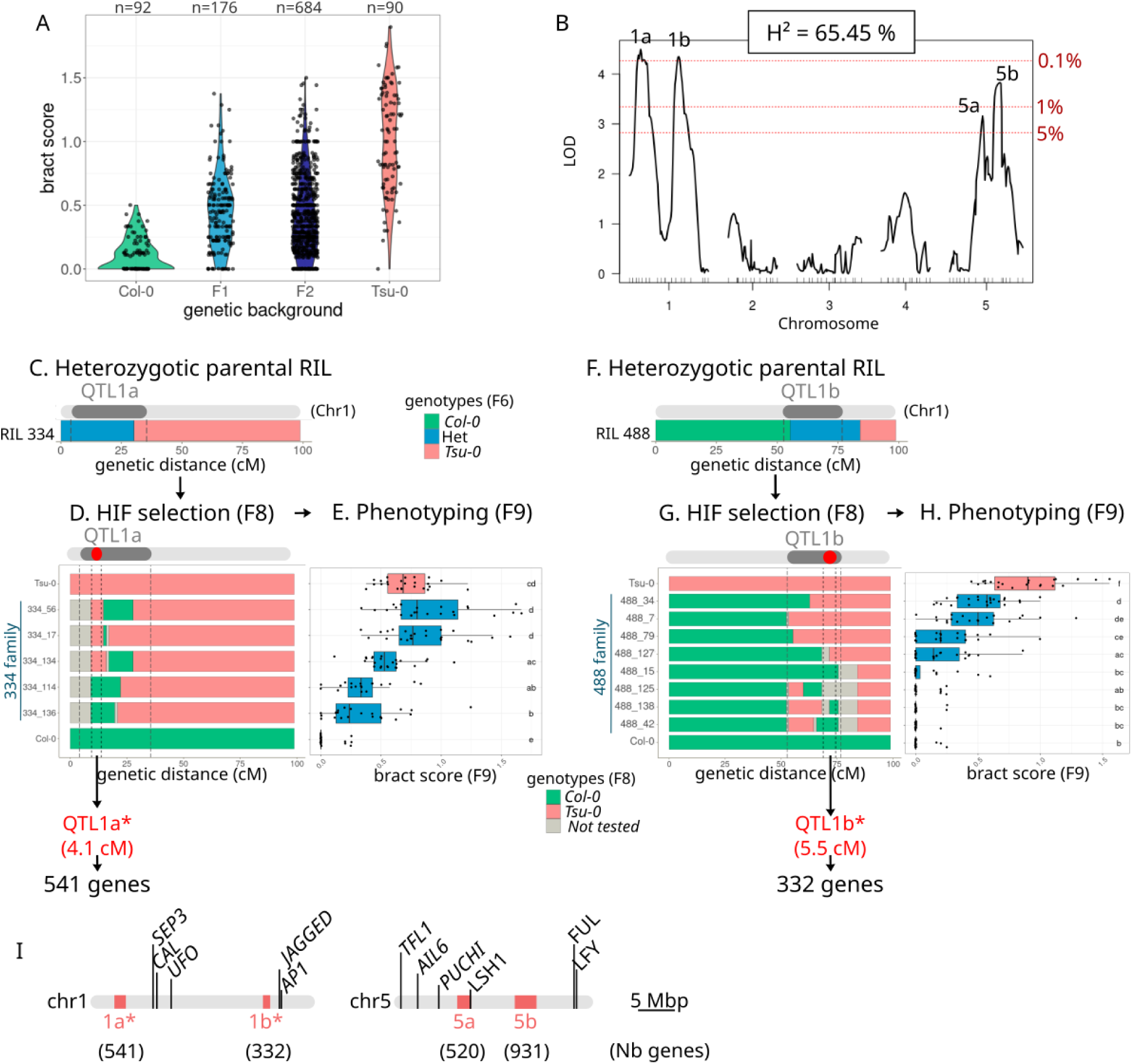
Basal bract formation in *Tsu-0* is controlled by four major QTL that do not contain the FMI genes associated with bract repression. **A**, Genetic transmission of bract score in F1 and F2 hybrids from a cross between *Col-0* and *Tsu-0* parent plants. **B**, QTL mapping for bract score in a set of RILs identifies four putative QTLs, two in chromosome 1 (1a and 1b) and two in chromosome 5 (5a and 5b). Dotted horizontal red lines indicate three different significance thresholds computed from 2000 permutations. On top of the graph, H^2^ indicates the (broad-sense) heritability of the bract score computed among the RIL set. **C**, Genotype of the F6 generation of the line RIL334 in chromosome 1, showing a heterozygotic region overlapping with QTL1a. **D**, Chromosome 1 genotypes of selected lines forming a heterologous inbred family (HIF) obtained from RIL334 after two more generations of selfing (F8). **E**, Box plots of bract scores of the different lines from the “334 family” (blue boxes) with parental controls (*Col-0*: green, *Tsu-0*: red, N > 20 plants per line). Segregating bract scores within the family map a narrower region, QTL1a*, spanning 541 annotated genes. **F**, Genotype of the F6 generation of the line RIL488 in chromosome 1, showing a heterozygotic region overlapping with QTL1b. **G**, Chromosome 1 genotypes of selected HIF lines obtained from RIL488 at F8 and (**H**) a box plot of their bract scores (blue boxes), with parental controls (*Col-0*: green, *Tsu-0*: red, N > 21 plants per line). This maps a narrower region, QTL1b*, spanning 332 annotated genes. In E and H, lines not sharing the same letter(s) are statistically different (*post hoc* Tukey analysis with 0.05 sign. level from a glm of bract scores fitted with a quasi-poisson distribution). **I**, Final genetic mapping of the ‘basal bract’ trait in *Tsu-0* and the location of genes reported to impact bract development in previous mutant studies.

To better estimate the QTLs, we also used Recombinant Inbred Lines (RIL), which allowed averaging of bract scores across isogenic plants. Using an existing RIL set (Simon et al. 2008), we measured the average bract scores of several lines (**Fig. S5A**) for which we checked the absence of recombination bias (**Fig. S5B**). The heritability of the bract score was high among RIL (65.45%

+/− 1.7) and a single QTL scanning mapped four peaks: two most significant on chromosome 1 (LOD score > 4 with a p-value < 0.1% named 1a and 1b) and two less significant on chromosome 5 (LOD score > 3, named 5a and 5b; **Fig. 3B**). Consistent with the BSA result, this analysis provided slightly different QTL positions and shorter intervals. A two-QTL scan suggested additive effects between QTLs 1a and 1b (**Fig. S5C**). To confirm and reduce the intervals of these two major QTLs, we used two RILs identified as heterozygous in a region overlapping with these QTLs at F6 generation (**Fig. 3C-H**). F_8_ plants from these RIL were re-genotyped at a higher density with new genetic markers (**Fig. S6A,B**) to select recombining intervals homozygous for either allele of the two parents (**Fig. 3D, G** and **Fig. S6F**). Selected plants were used to generate new lines forming a heterologous inbred family (HIF) with identical genotypes except in a small interval, which allows testing the effect of alleles in that region only (see **Fig. 3D**). Scoring bract among HIF confirmed both QTLs and mapped them to smaller intervals named 1a* and 1b*.

Additive effects of QTL 1a and 1b appeared when *Tsu-0* allele in 1b* was combined with *Col-0* allele in 1a*, the bract score reaching about half of the *Tsu-0* parent bract score (see in 488 families, **Fig. 3G, H**). However, *Tsu-0* parent score is tied only when both QTLs bear *Tsu-0* alleles (see in 334 families, **Fig. 3D, E**). As a control, we generated a HIF to test the region between 1a and 1b and demonstrated that it did not influence the bract score (**Fig. S6C-E**).

The final mapped intervals remained large (4.1 to 11.8 cM), containing many genes (332 to 931, 2324 annotated genes for the four QTL, **Fig. 3I** and **Table S2**). The parental accessions that we re-sequenced differ by over 770,000 small genomic variations (SNPs and indels), with more than one variation every 175 bp on average, leaving few invariant genes to exclude (**Table S2**). However, known bract-related genes were absent from the QTLs (**Fig. 3I**), whether those genes were reported as promoting bract development (e.g. *JAGGED*) or as being involved in the FMI pathway and acting as bract repressor genes (**Table S1**). Only *LSH1* was found to lie at the border outside QTL 5a. This absence of major FMI genes in our QTL is consistent with the wild-type phenotype of flowers associated with bracts in *Tsu-0* (**Fig. 1G**).

To decipher the causal variation responsible for basal bracts, we took advantage of pleiotropic phenotypes co-segregating with basal bracts in the RIL. The number of cauline branches positively correlated with the bract score and QTL1b overlaps with a QTL controlling cauline branch number (**Fig. S7**). This suggests that the gene controlling basal bracts in QTL 1b* might also promote more cauline branches. Second, “branched siliques” could sometimes be observed among RILs and HIF at low penetrance (**Fig. S8A,B**). We interpreted this transgressive phenotype as resulting from an incomplete floral determinism with reiteration of floral bud formation in whorls, as reported for mutants such as *ap1* (Irish and Sussex 1990). In HIF 488, this phenotype occurred only when *Tsu-0* QTL 1b* was associated with *Col-0* haplotypes in other mapped QTLs (**Fig. S8C**). In HIF 334 however, this phenotype was not correlated to a consistent genetic combination of mapped QTLs (**Fig. S8C**). This transgression suggests that the maintenance of a wild-type phenotype in *Tsu-0* bracteate flowers relies on genetic interactions and may interfere with the FMI pathway.

To conclude, our genetic analysis demonstrates that the genetic control of basal bracts in *Tsu-0* is complex, involving several loci with additive effects on bract development and epistatic interactions with other traits. It also suggests that the causal genes differ from those previously identified in mutants with bracts.

### Time-series transcriptomics reveal a peak of gene expression divergence at the floral transition, while FMI genes remain synchronized

Developmental programs are governed by gene regulatory networks, often leading to substantial transcriptional changes. We therefore sought to identify genes whose expression changes with bract development. We profiled meristem transcriptomes before, during and after floral transition in both accessions. Microdissected meristems were matched to the same four developmental stages: vegetative, late vegetative, transition and floral (V, L, T and F, respectively, **Fig. 4A**). We performed RNAseq and used principal component analysis (PCA) to assess the consistency between replicates (**Fig. S9B**).

**Figure 4:**
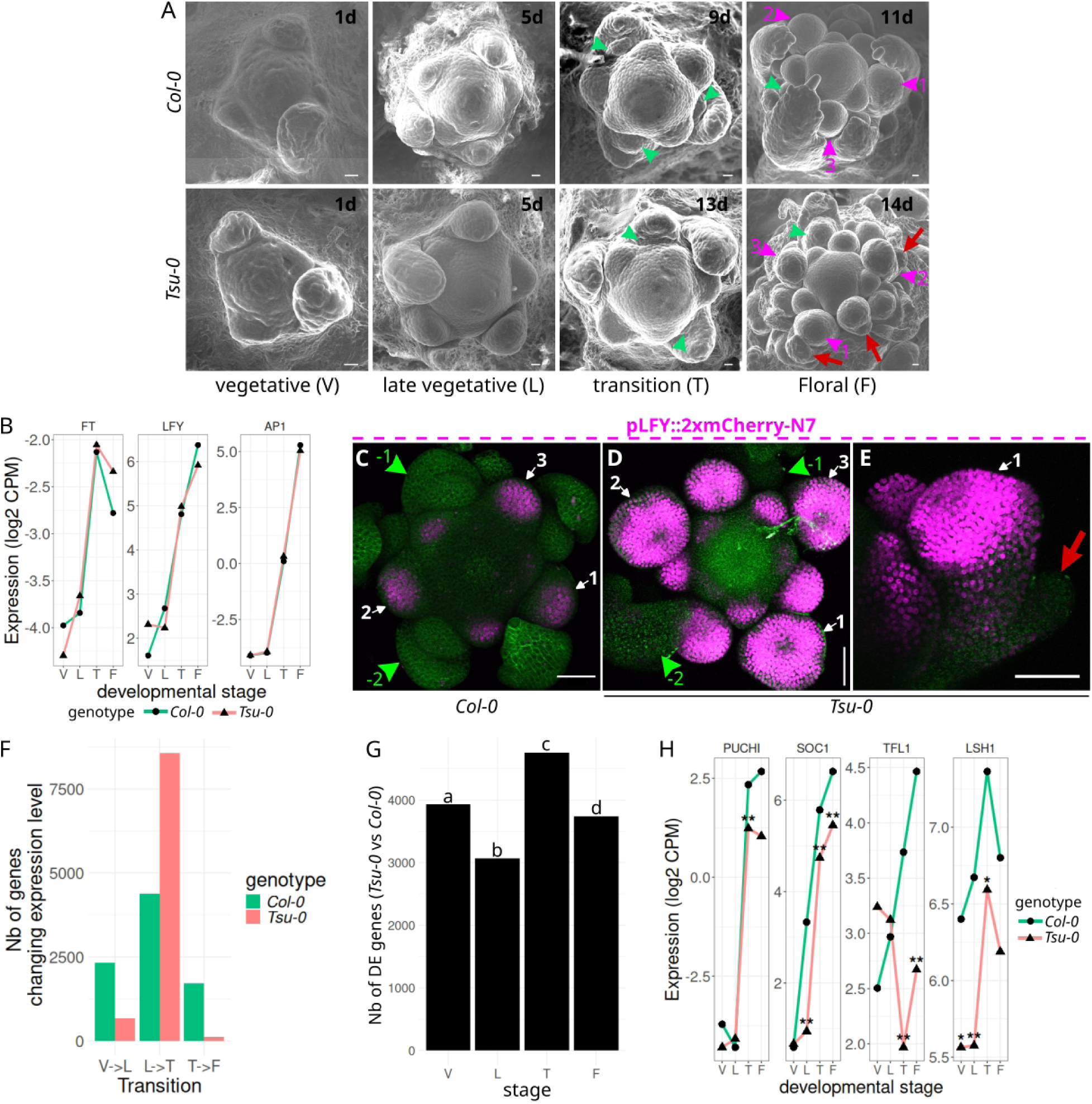
Transcriptomic divergence between *Tsu-0* and *Col-0* meristems peaks at the floral transition despite synchronization of the FMI pathway genes. **A,** Scanning electron microscopy showing the evolution of the main meristem in both *Col-0* and *Tsu-0* at the four different stages (V, L, T, F) used for RNAseq. Plants were first synchronized by 21 days of a non-inductive short-day light regime before a transfer to inductive long days. The date after the transfer is indicated in the top-right corner of each picture. Green arrowheads point to branches (with a leaf) while magenta arrowheads point to the first flowers produced after floral transition. In *Tsu-0*, red arrows show bracts. Scale bars: 50 µm. **B,** Dynamics of the expression levels of FT, LFY and AP1 (three important regulators of floral transition and identity) in both *Col-0* (green) and *Tsu-0* (red) over the four developmental stages. Differential expression analysis reveals no difference at any stage between the two accessions. **C, D**. Confocal live imaging of the main meristems of *Col-0* (**C**) and *Tsu-0* (**D**) plants at floral transition, expressing a pLFY transcriptional reporter (magenta) and a membrane marker (green) for morphology (top-view projections of stack acquisitions). Green arrowheads: branches, ordered with decreasing numbers from the floral transition; white arrows: flowers ordered with increasing numbers from the floral transition. In both genotypes, pLFY expression is absent from branch meristems and suddenly appears in the first flower onwards. Representative pictures of at least 6 plants per genotype captured at floral transition. **E**, side-view of the first flower from image D (*Tsu-0*): a bract (red arrow) is visible on the abaxial side of this young flower. Scale bars: 50 µm. **F**, Bar plots of the number of genes differentially expressed between two consecutive stages in *Col-0* (green) and *Tsu-0* (red). The number of changes peaks at the V-to-T stage transition, especially in *Tsu-0*. **G**, Bar plots of the number of differentially expressed genes (DEG) between *Col-0* and *Tsu-0* at each stage of the time course. T stage is when the highest number of DEG is measured (on top of the bar, different letters indicate the statistical difference with a chi-square test of homogeneity after *post hoc* analysis). **H**, Dynamics of the expression levels of four genes reported to be involved in bract development, showing a significant difference at least at the T stage between *Col-0* (green) and *Tsu-0* (red). One star: genes are differentially expressed; two stars: the absolute fold change is superior to 1.

To monitor the progression from vegetative growth to flowering at the transcript level, we analysed the expression dynamics of several key regulators of floral transition, such as *FLOWERING LOCUS T (FT)* (Wigge et al. 2005), and of flower identity (e.g. *LFY* or *AP1).* The synchronized expression of these genes in the two accessions validated the precision of our morphological staging (**Fig. 4B**): despite different absolute flowering times (**Fig. 4A**; **Fig. S3**; **Fig. S9A**), our experimental design realigned the two accessions’ developmental progression using the floral transition as a common timer. Hence, gene expressions that desynchronize from flower formation can easily be tracked. To ensure that our bulk RNAseq data does not obscure changes in gene expression patterns related to flower identity, we introduced an LFY transcriptional reporter in both accessions to precisely monitor the activation of this master floral identity gene in space and time during the floral transition. Live confocal imaging revealed similar dynamics in both genotypes, with no expression of *LFY* in branches and high expression levels in all flowers from the first flower onwards (**Fig. 4C**), regardless of bract presence in *Tsu-0* (**Fig. 4D,E**). These results, in line with our genetic experiments, confirm that alteration of *LFY* dynamics during floral transition is likely not involved in the delayed repression of bracts. As previously reported (Chahtane et al. 2018), bracts in *lfy* mutants are not precisely located at the base of branches (**Fig. S10**), as opposed to *Tsu-0* basal bracts (**Fig. 1B**). This indicates that the developmental contexts leading to the formation of bract in mutants versus basal bracts in natural accessions may be really distinct, in line with the involvement of distinct genetic pathways shown above (**Fig. 3**).

In the time-course, the stage T should be the most relevant to study bract formation even if no bract is visible at this stage. Indeed, bract and flower initiations start before any morphological event (Heisler et al. 2005). The delay between stages T and F (a median of 1 to 2 days, **Fig. S9A**) is compatible with the initiation of the first flowers and their bracts at stage T. Hence, transcriptional changes associated with the delay of bract repression should transiently appear at stage T and progressively fade by stage F (since only the 1 to 5 first flowers present a bract, **Fig. 1**). At the global transcriptome scale, the stage T stood out as a critical transcriptomic transition in both accessions, with the highest number of gene expression changes upon reaching this stage (**Fig. 4F**). Interestingly, the divergence between the transcriptomes of the two accessions (as measured by the number of differentially expressed genes -DEG) also peaks at stage T, with up to 4,759 genes DEG (**Fig. 4G, table S3**), including genes potentially responsible for bract formation. In conclusion, the relevant time window during which bract repression is uncoupled from flower development corresponds to a period of highly dynamic gene expression, both throughout a plant’s development and among plants with different genetic backgrounds.

### Transcriptomic data identify candidate genes responsible for the desynchronization of bract repression

To identify putative genes impacting bract formation among DEGs at stage T, we first cross-referenced our transcriptomic (4,759 DE genes) and QTL-mapping data (2324 genes in intervals). We further filtered out genes that had no genetic polymorphisms (including variations with no putative functional impact, see methods) between *Col-0* and *Tsu-0*. This selection yielded a list of 362 candidate genes spread over the four QTLs (**Table S4**), including 24 transcription factors, and none of them had previously been associated with bract formation. This suggests that basal bract formation in a natural accession can be genetically determined by a novel set of genes likely present in this list.

To determine whether these causal genetic variations influence the expression of genes known for their role in bract development (**Table S1**) or act through an independent mechanism, we systematically analyzed the expression dynamics of these genes. Only *PUCHI*, *SOC1*, *TFL1* and *LSH1* significantly differed at stage T between *Tsu-0* and *Col-0* (**Fig. 4H** and **Fig. S9D**). Single *puchi*, *soc1* or *lsh1* null mutations alone do not induce basal bracts (Rieu et al. 2024; **Fig. S10C,D**), so the isolated effect of their reduced expression in *Tsu-0* at stage T (54%, 52% and 41%, respectively) does not likely impact bract development. By transforming lateral branches into flowers, *tfl1* mutants display basal bracts (**Fig. S10C,D**). But *tfl1* phenotype is recessive: no basal bract exists on heterozygous TFL1//tfl1 plants (data not shown), so the effect of the reduced *TFL1* expression at stage T in *Tsu-0* (71%) is again uncertain. To our knowledge however, combinations of these mutations have not been reported. Moreover, these gene expression differences persisted in stage F while bracts disappeared with older flowers: hence, the reduced expression of *PUCHI*, *SOC1*, *LSH1* and *TFL1* is not correlated to bract formation at all developmental stages. Altogether, this suggests that both the natural genetic determinism of delayed bract repression and the underlying transcriptional program acting in the meristem are largely independent of the genetic pathway identified in bract mutants.

### Using time-series to identify a transcriptomic signature of leaf/bract development in shoot meristem

To identify biological functions specific to the critical stage T in *Tsu-0* without prior assumptions, we performed a GO term analysis on all DEGs compared with *Col-0*. However, the results were not very informative, notably lacking terms related to leaf development (**Fig. S11A,C**). Since *Tsu-0* and *Col-0* are distant backgrounds, it is not surprising that many gene expression differences at stage T are not related to bract formation or even to development (see the second axis of PCA, **Fig. S9B)**. To specifically isolate the gene regulatory network responsible for delayed bract repression in *Tsu-0*, and thanks to the information provided by the time-series, we designed a clustering method to isolate a transcriptomic signature common to all leaf-producing stages versus non-leaf producing stages. Briefly, expression data were gathered in two groups corresponding to either bract-less (*Col-0* T, F, and *Tsu-0* F) or leaf/bract-producing meristems (the other samples, including *Tsu-0* T) and we selected genes whose expression profile clusters these two groups apart (**Fig. 5C**). These genes can be considered “bract signature genes”, with a positive or inverse correlation with the presence of bract if their expression is up- or down-regulated (respectively) at stage T in *Tsu-0*.

**Figure 5:**
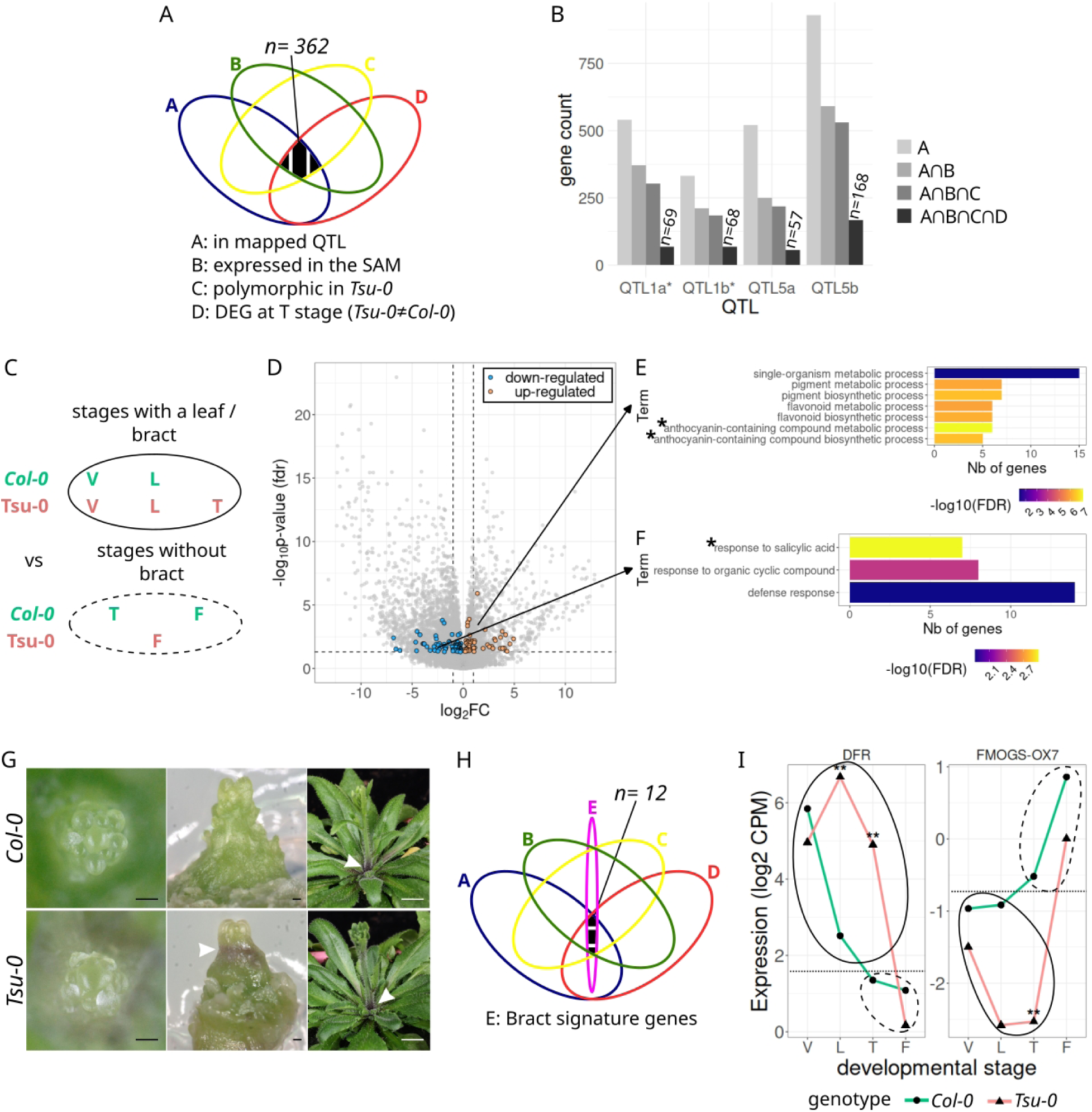
Identifying a transcriptomic signature of leaf development in shoot meristems at the floral transition stage using time-series RNAseq. **A**, Intersecting genetic mapping and transcriptomic data: if a genetic variant causing *Tsu-0* bracts also induces the differential expression of the gene in *cis*, this gene should verify the conditions defined by A, B, C and D. **B,** evolution of the number of candidate genes mapped in the four QTL (condition A) by the successive addition of other conditions (B to D). **C**, Strategy to cluster genes based on the presence (circle with the plain line) or the absence (dotted circle) of leaf and/or bract in the different combinations of stages and genotypes (see text for details). **D**, Volcano plot of gene expressions at the stage T between *Tsu-0* and *Col-0*. All genes expressed in the SAM are plotted (n=21,652 grey dots) but only the genes fulfilling the clustering condition defined in C (“bract signature genes”) are highlighted in orange and blue for up- and down-regulation, respectively (n=124). Vertical dashed lines: absolute fold change superior to 1, horizontal dashed line: significance threshold at 5.10^−2^ (adjusted p.value with fdr method). **E**, Seven ‘biological process’ GO terms are statistically enriched among the up-regulated genes identified in D (bar plot providing the number of related genes). They are all related to the anthocyanin biosynthesis pathway, which is cited in the two most significant terms (stars). **F**, Only three ‘biological process’ GO terms are statistically enriched among the down-regulated genes identified in D (bar plot providing the number of related genes); they collectively point to defense response and more specifically the acid salicylic pathway (star). In E,F, color gradients indicate the statistical significance of the GO term enrichment (Fisher’s test with Yekutieli adjustment method). **G**, Representative pictures of micro-dissected meristems in *Col-0* (upper row) and *Tsu-0* (lower row) just at or before (left) or after (middle) stage T (N>15 for each genotype) and a close-up of the base of the bolted main stem (right). At stage T, *Tsu-0* meristems display a typical anthocyanin red coloration just below the meristem, which is not observed in *Col-0*. After bolting, both genotypes show anthocyanin coloration at the base of the stem. Scale bars: 100 µm (left and middle), 1 cm (right). **H**, Intersection of the gene set in A with the ‘bract signature genes’. **I**, Examples of the expression profile of two candidate genes, showing an up- (left) and a down- (right) regulation at the T stage. *DFR* is an enzyme involved in anthocyanin biosynthesis while *FMOGS-OX7* is annotated as a salicylic acid responding enzyme. Two stars mean that the genes are differentially expressed and the absolute fold change is superior to 1. The bract and bract-less clusters are outlined with a solid and dashed circle, respectively, while the horizontal dotted line highlights their separation.

Only 124 genes met these criteria (**Fig. 5D, Table S5**), including *SOC1*, whose expression anti-correlates with the bract presence. Interestingly, two enriched GO-terms emerged from this list of bract signature genes: anthocyanin biosynthesis and salicylic acid (SA), associated with up- and down-regulated genes, respectively (**Fig. 5E**). Backtracking these ontology terms for all the 4,759 DEGs at stage T (**Table S6** and **S7**) retrieved even more genes associated with these pathways, supporting that their activity levels differ between *Tsu-0* and *Col-0* at the critical stage T (**Fig. S11D**). High anthocyanin production in *Tsu-0* at stages L and T was evident from the frequent purple coloration just below the meristems, contrasting with the green tissues in *Col-0* (**Fig. 5G**). Sometimes, this purple color extended to young organs where bracts were initiated (**Fig. S11E**). As the stem grew, the pigment receded to the rosette junction in both accessions (**Fig. 5G**), vanishing from the apical meristem. This trait specific to *Tsu-0* validates our clustering strategy: it is able to capture genes whose expression differ between *Tsu-0* and *Col-0* at stage T and with overall contrasted levels between vegetative and flower stages, as expected for a gene regulating bract repression in synchronization with flower development.

Interestingly, among these 124 bract signature genes, 12 also lie in the mapped QTLs (**Fig. 5H**), including one anthocyanin biosynthetic enzyme (the dihydroflavonol reductase, DFR) and five SA-responding genes (see **Table S5**). For example, **Figure 5I** shows the expression profiles of two such genes. However, further work is required to test whether these candidates are responsible for basal bract formation in *Tsu-0* and whether the anthocyanin and/or SA pathways are involved in this natural variation.

### Curve registration revealed massive gene desynchronization and challenged the hypothesis of a “prolonged vegetative state” during floral transition in *Tsu-0*

In the previous clustering approach, both up- and down-regulated genes display similar temporal dynamics: they tend to be delayed in *Tsu-0* compared to *Col-0* (**Fig. 5I**). This result suggests that a transcriptomic leaf signature is maintained at the stage T in *Tsu-0* compared to *Col-0.* This desynchronization of the expression of some genes with the developmental clock imposed in our experiment (from V to F stage) identifies transcriptomic heterochronies, i.e. changes in the timing of gene expression.

To assess the importance of heterochrony in the divergence of *Tsu-0* and *Col-0* transcriptomes, we first tested whether its effects could be detected in a global unsupervised analysis of gene expression levels over the time-course. We therefore performed a PCA and plotted the two main axes of variance after grouping samples by genotype and time points (**Fig. 6A**). While the second axis of variance (∼24.4%) differentiates genotypes, the main axis (47.9% of variance) orders the sampling time points chronologically in both genotypes, representing a transcriptomic age. Surprisingly, *Tsu-0* stage T does not align with *Col-0* stage T on this axis: it is closer to a stage F, explaining the limited expression changes when progressing to the next stage F (**Fig. 4F**).

**Figure 6:**
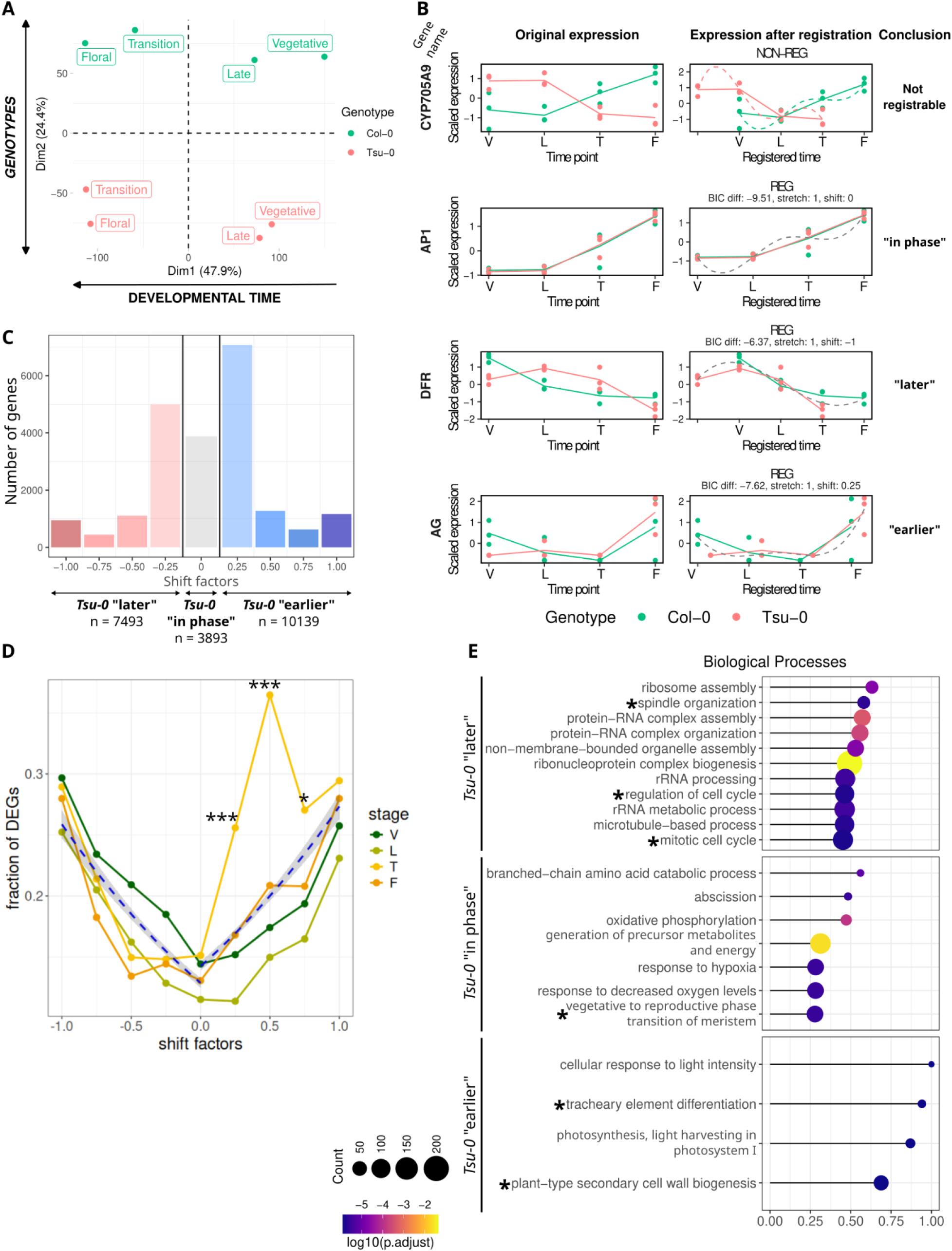
Massive and complex transcriptional desynchronisation coexist in *Tsu-0* across the floral transition. **A**, PCA of RNAseq data from microdissected meristems of *Col-0* and *Tsu-0* over the four sampled stages (biological replicates averaged per time points). The two main axes of the PCA can be interpreted as developmental time and genotype, respectively. **B**, Examples of temporal registration of gene dynamics (right panels) between *Col-0* (green) and *Tsu-0* (red) from scaled expression levels (left panel). Dots represent the expression levels of independent biological replicates, while lines indicate the mean expression level at each time point. In the right panel, the green (or red) dotted curves represent the fitted models for *Col-0* (or *Tsu-0*) independently, while the grey dotted curve represents the joint model for both *Col-0* and *Tsu-0*. If the green and red dotted curves are used, it means the two time series are best explained by two independent models, indicating they are not similar. Conversely, if the grey dotted curve is used, it suggests that a single model best explains both time series, indicating they are similar. The name of the gene plotted is indicated on the left of each row. These four genes exemplify the four possible categories (from top to bottom, respectively): genes that cannot be registered (N = 43, *e.g. CYP705A9*), genes with identical temporal dynamics (shift = 0, *e.g. AP1,* see also Fig. 4B) and genes whose expression dynamics in *Tsu-0* must be shifted negatively (*e.g. DFR,* see also Fig. 5I) or positively (*e.g*. *AG*) to align with *Col-0*. The last column provides a biological interpretation of the computed shift (see main text): a null shift indicates that the expression dynamics in *Tsu-0* stay “*in phase*” with floral transition while negative or positive shifts indicate that the expression dynamics in *Tsu-0* are desynchronized later or earlier, respectively, than the phenotypic progression of floral transition. **C**, Distribution of heterochronic shifts between *Tsu-0* and *Col-0* on the entire meristematic transcriptome, computed by the registration method from B. The shift value is colour-coded in a red-to-blue gradient from −1 to 1. **D**, Correlation between the heterochronic shifts in gene expression and the fraction of genes presenting a differential expression between *Tsu-0* and *Col-0* for each stage, colour-coded from green to orange. The blue dotted lines are two piecewise linear regressions modeling the effect of the shift with a binomial model for positive and negative values (p-value < 2e-16) surrounded by a 5% confidence interval (gray shade). Stars highlight the significantly higher proportion of DEGs at the T stage (p-value code: *** < 0.001, * < 0.05; see also figure S12 for the complete statistical analysis). **E**, GO term enrichment analysis associated with the three categories of heterochronic shifts. The list of all significant ‘biological process’ GO terms (BH-adjusted p.value < 0.05) was simplified using semantic similarity (cutoff = 0.7) and the Rich Factor was computed for the remaining terms, revealing the proportion of genes involved among all the genes associated with this GO term. Dot size indicates the count of genes and the color scale is the statistical significance (BH-adjusted p.value) of the enrichment in the shift category. Stars indicate GO terms referring to developmental processes.

As suggested by Calderwood et al. (2021), the transcriptomes of two genotypes during flowering cannot be aligned to a single developmental time; each gene may desynchronize differently, sometimes in opposite directions. To quantify each gene desynchronization between *Col-0* and *Tsu-0*, we used curve registration (Calderwood et al. 2021) that can predict subtle temporal shifts of gene expression (Kristianingsih, 2024). We analyzed the shift relative to the floral transition, used as a common reference clock between the two accessions. Thus, genes with no shift, like *AP1* (**Fig. 4B and Fig. 6B**), can be nevertheless shifted in absolute time, because *Tsu-0* plants flower later with an older absolute age (**Fig. S3 and S9A**). In our approach, positive and negative shifts reflect desynchronization relative to the floral transition and gene expression dynamics may shift earlier or later. Unlike our previous clustering approach (**Fig. 5C**), this registration method uses a common scaled expression level and computes only temporal shifts, regardless of absolute expression levels (**Fig. 6B** and **Fig. S12**).

Only a few genes could not be registered (**Fig. 6B**, N = 43), suggesting that most genes follow similar temporal dynamics in both accessions (see examples **Fig. S9C, D** or **Fig. 4B, H**). Most genes could be classified into three categories: null, negative and positive shifts (**Fig. 6B**). A null shift indicates that the gene expression in *Tsu-0* stays “in phase” with flowering, like *AP1* (compared with **Fig. 4B**). A positive or negative shift indicates that gene dynamics in *Tsu-0* occur earlier *(e.g. AG*) or later (*e.g*. *DFR*), relative to the floral transition, revealing heterochronies between transcriptomic and phenotypic processes. Transcriptome-wide, the shifts were broadly distributed, showing massive and complex desynchronisation of transcriptome dynamics (**Fig. 6C**). Hence, the floral transition in *Tsu-0* did not impose a synchronization to the entire meristematic transcriptome: only ∼18% of genes (n=3879) stayed in phase with this process while the majority showed delayed or advanced expression dynamics. Even more genes shifted earlier relative to the floral transition, supporting the PCA interpretation (**Fig. 6A**). Since curve registration computes temporal shifts in gene expression regardless of absolute expression levels, we investigated whether these shifts could affect differences in gene expression levels. Indeed, there is a strong correlation (p-value < 2e-16) between the absolute shift value and the proportion of DEGs between *Tsu-0* and *Col-0* at all developmental stages (**Fig. 6D**). This suggests that the desynchronization of gene dynamics between the two accessions contributes to differences in gene expression levels. Furthermore, this effect is significantly higher at the T stage for positive shift values ranging from 0.25 to 0.75 (**Fig. 6D** and **Fig. S12**): this suggests that genes with “earlier” dynamics in *Tsu-0* explain at least partially the peak of transcriptome divergence observed at the T stage (**Fig. 4G**).

In conclusion, the differences in the transcriptomes of *Tsu-0* and *Col-0* are impacted by a significant level of heterochrony, with complex combinations of temporal shifts. Although the bract is often considered a ‘juvenile’ vegetative structure, its formation at the floral transition does not indicate a global juvenilization at the transcriptomic level.

### Investigating which biological processes are affected by desynchronization of gene expressions in meristems during flowering

As expected, genes in phase were primarily linked to flowering and developmental phase change as shown by key regulators of these pathways (**Fig. S12B, C**), which also emerges from GO term enrichment analysis (**Fig. 6D**). In contrast, genes previously associated with bract mutants exhibit a wide range of shifts, from very early to very late. This shows that they are variably affected by these desynchronizations (**Fig. S12A, C**), reinforcing the idea that they are not involved in basal bract formation. GO term analysis for each shift category highlighted the particular processes desynchronized from flowering in *Tsu-0* compared to *Col-0* (**Fig. 6D**). For instance, vascular differentiation (two GO terms mentioning tracheary element and secondary cell wall) emerges as a process shifted earlier in *Tsu-0* relative to the floral transition (**Fig. 6D**). Vascular differentiation dynamics may be driven by other progression factors like absolute age, suggesting a loose connection with the flowering pathway. Conversely, processes shifted later in *Tsu-0* revealed terms related to cell division (spindle, cell cycle, mitosis) and ribosomal biogenesis (six GO terms mentioning ribosome, rRNA, ribonucleoprotein and protein-RNA complexes; **Fig. 6D**), which suggests that core meristematic functions are prolonged in *Tsu-0*. Further work is yet needed to explore whether and how the desynchronization of these processes relative to floral transition impacts bract de-repression.

## Discussion

In this study, we investigated the transient formation of ‘basal bracts’ at the base of inflorescences in natural accessions of the model plant in *A. thaliana*. We characterised basal bract formation at developmental, genetic and transcriptomic levels. We found that this phenotype is associated with a desynchronization of bract repression from flower formation during the floral transition.

### The mechanisms desynchronizing bract repression and flower development

Evidence from the literature suggests that flower development, through redundant actions of the genes from the FMI pathway, represses bract development, hence naturally coupling flower establishment and bract repression. However, natural basal bract demonstrate that these two processes can be uncoupled in a short time window during the floral transition when both flowers and bracts develop together.

The wild-type phenotype of basal bract associated flowers (**Fig. 1**) and the proper dynamics of many FMI genes (e.g. *LFY*, *AP1,* **Fig. 4**) suggest that the transient bract de-repression at floral transition is not a consequence of an impaired FMI establishment, contrary to previously reported mutants. In line with this, bracts are rarely produced with the first flowers in mutants of the FMI pathway (**Fig. S10**). Our quantitative genetic (**Fig. 3**) and transcriptomic data (**Fig. 5**) further show that the genes causing this transient bract de-repression are different from the handful of FMI genes whose mutants display bracts.

However, the precise mechanisms involved remain unresolved. Finer mapping or alternative quantitative genetic approach such as GWAS could help pinpoint causal genes for their definite validation. However, the four identified QTLs, including two with additive effects (**Fig. 4 and Fig. S5**), indicate that several complex genetic mechanisms may be at work. Our transcriptomic analysis established correlations of differentially expressed genes and pathways in the meristem at the time when bract de-repression occurs: up-regulation of anthocyanin pathway, down-regulation of salicylic acid pathway (**Fig. 5**) or prolonged activity of ribosome biogenesis or cell cycle (**Fig. 6**). This however, does not demonstrate their role in promoting bract development: a transcriptomic signature of leaf development in stage T meristems could simply be a consequence of the developing bracts rather than its cause.

Interestingly, several plant species evolved leaf-less inflorescences, such as grasses (Poaceae), through apparently convergent and unrelated mechanisms (Mach 2010). Indeed, *Arabidopsis* and *Poaceae* mutant bracts are not strictly homologous and so far, different genetic mechanisms were reported (Whipple et al. 2010; Whipple 2017; Xiao et al. 2022). However, some of these mechanisms are still partially unresolved (Kawakatsu et al. 2009; Wang et al. 2021). Since in both groups, bract exclusion from wild-type inflorescences involve a leaf repression mechanism synchronized with a developmental phase change, further research on these two groups may reveal whether they share more similarities than what is currently described.

### Linking floral transition, transcriptome divergence and developmental robustness

We found that gene expression differs the most between the two accessions at the floral transition, at the time when bracts are de-repressed in *Tsu-0* (**Fig. 4**). Furthermore, curve registration identified an important level of gene desynchronization, which likely contributes to the divergence in gene expression levels, especially at the stage T (**Fig. 6**). Specifically, these desynchronizations alter the expression timing of many genes relative to a minor gene pool driving the floral transition, which we used as a reference in our experiments.

As flowering is a key adaptive trait that involves transcriptional reprogramming, the floral transition may be a developmental stage that leads to large transcriptome differences among natural populations. Indeed, massive and complex gene desynchronisation during floral transition were reported at both intra- and inter-species level in *Brassica rapa* cultivars and *A. thaliana* (Calderwood et al. 2021). This phenomenon may arise from the strong selection pressure adapting flowering time to the environment (Bloomer and Dean 2017). As a result, the flowering-related genes could shift and desynchronize from numerous others not functionally related to flowering, creating a peak of transcriptome divergence in the short time window when flowering-related genes vary quickly (**Fig. S13**). In line with this, a peak of gene expression divergence has also been reported in meristems transitioning from vegetative to floral stages between different species of Solanaceae (Lemmon et al. 2016).

In different unrelated phyla like animals, plants, brown algae or fungi, the “hourglass” (Kalinka et al. 2010; Quint et al. 2012; Cheng et al. 2015; Lotharukpong et al. 2024) and “inverse hourglass” (Lemmon et al. 2016; Levin et al. 2016) models propose that reduced, or respectively increased, transcriptome divergence at specific stages of development drive lower (respectively higher) phenotypic variations in the final body plans. For instance, the high transcriptome divergence between *Solanaceae* species during floral transition correlates with the diversification of their inflorescence architectures (Lemmon et al. 2016). Here, we report a similar correlation within natural populations of *A. thaliana*: the transcriptome divergence peaks at floral transition, which correlates with a developmental variation, the formation of basal bracts. Interestingly, several observations suggest that developmental robustness (i.e. the ability to produce an invariant phenotype) is reduced during floral transition in *Brassicaceae*, at both inter- and intra-species levels. In addition to basal bracts (**Fig. S1**), botanists have also documented examples of frequent flower-to-shoot transformations or flower dimorphisms (Arber 1931a; Arber 1931b). A possible explanation would be that, as a consequence of the constant adaptation of flowering time, the many changes in gene expression lead to desynchronisation of genetic programs occurring during the short floral transition, creating new, transient gene expression states that fuel developmental variation (**Fig. S13**). More generally, this questions whether rapid developmental transitions of adaptive traits that change the timings of events, may result in more gene desynchronization and developmental variation. To test this hypothesis for the case of bract development, further investigation into the identification of the genetic and developmental mechanisms that uncouple bract repression from flower formation in natural *A. thaliana* accessions will be important. Such follow-up studies may also shed light on how the presence of bracts has frequently evolved within and between species, both at the base of branches in link with floral transition and in entire inflorescences as well.

## Materials and Methods

### Plant growth conditions

Seeds were sown on peaty-clay soil, stratified at 4 °C for at least two days, and watered with fertiliser (18-10-18 N-P-K) under LED lighting (sunlight spectrum NS12, 150 µmol.m-2.s-1). Three different day/night regimes were used in the experiments: short-days (SD) with 8 h light and 16 h dark (non-inductive photo-conditions for flowering); long-days (LD) with 16 h light and 8 h dark and continuous light (CL) with 24 h light (both photo-inductive flowering conditions). Temperature and humidity are controlled as follows: 22 °C and 60% humidity during light for LD and CL conditions, and 18 °C and 70% humidity constantly in SD and during night time in LD. For the Bulk Segregant Analysis and the RNAseq time course, plants were grown 20 days in SD before switching to LD. For the RIL phenotyping assays, plants have been directly cultured in LD.

### Plant materials

*Arabidopsis thaliana* natural accessions and associated RILs were obtained from the Versailles *Arabidopsis* Stock Centre (VASC), *Col-0*: 186AV, *Tsu-0*: 91AV, RIL set name: 3RV. F_1_ and F_2_ plants used for the BSA were generated by crossing the parents *Tsu-0* and *Col-0* in both directions. The following strains were obtained from the NASC and are in *Col-0* background if not otherwise mentioned: *tfl1-13*: N6237; *tfl1-14*: N6238. *puchi-1, bop1-4* x *bop2-11* and *puchi-1 x bop1-4* x *bop2-11* (Karim et al. 2009), *lfy-12* (Maizel and Weigel 2004) and *soc1-2* (Michaels et al. 2003) were described previously. Plants expressing pLFY::2xmCherry-N7; 35S::Lti6b-YFP (in *Col-0* and *Tsu-0*) were generated for this study. *Lobularia maritima* and *Allaria petiolata* plants were natural specimens collected outside the laboratory, in France. The phylogeny of *Brassicaceae tribes* used in Fig. S1 is extracted from the Brassibase website (Kiefer et al. 2014).

### Plasmid constructions and plant transformation

<u>pLFY::2xmCherry-N7</u>: since no polymorphism was sequenced in the *LFY* promoter between *Col-0* and *Tsu-0* accessions, a sequence starting at −2277pb upstream of the ATG was amplified by PCR from *Col-0* genomic DNA using 5’GGGGACAACTTTGTATAGAAAAGTTGATCCATTTTTCGCAAAGG and 5’GGGGACTGCTTTTTTGTACAAACTTGAATCTATTTTTCTCTCTCTCTCTATC primers. PCR fragments were purified and inserted into pDONR P4-P1R with a gateway BP reaction. This plasmid was then recombined in a three-fragment gateway reaction with 2xmCherry pDONR211 and N7-tag pDONR P2R-P3 (containing the nuclear tag N7 and a stop codon) and the destination vector pK7m34GW. <u>35S::Lti6b-YFP</u>: a pDONR P2R-P3 plasmid containing the sequence of the plasma-membrane protein Lti6b (Cutler et al. 2000) was recombined in a three-fragment gateway reaction with a 35S pDONR P4-P1R and the YFP pDONR P2R-P3 into the destination vector pB7m34GW. Both constructs were then transformed into *Agrobacterium tumefaciens* C58pMP90 strain by electroporation and transformed into both *Col-0* and *Tsu-0* plants by floral dipping (Clough and Bent 1998). For each construct-by-genotype combination, several independent transgenic lines were selected in T_2_ for a single insertion event based on 3:1 resistant:sensitive mendelian segregation of the resistance provided by the transgene. The expression patterns of pLFY were reproducible between selected lines and matched published data for *Col-0*. We were unable to get a 35S::Lti6b-YFP line with a membrane signal as strong in *Tsu-0* as in *Col-0* (Fig. 2 and S2). However, despite the weak YFP signal, the morphology of the tissue could still be captured.

### Microscopic meristem imaging and image analysis

Meristems were imaged using a Scanning Electron Microscope (Hirox 3000 SEM), or with a confocal microscope Zeiss 700 LSM, according to the manufacturer’s instruction and without prior fixation. Multitrack sequential acquisitions were performed as follows: YFP, excitation wavelength (ex): 488 nm, emission wavelength (em): 300–590 nm; mCherry, ex: 555 nm, em: 300–620 nm. Detection wavelengths and laser power were identical for *Col-0* and *Tsu-0*, PMT voltage was identical for mCherry to allow comparisons of pLFY signal intensity in the two genotypes. The YFP PMT voltage was optimised on each plant. Confocal images were processed using imageJ (Schneider et al. 2012): maximum projections of mCherry channel and 16-bit-transformed standard deviation projections of the YFP channel were merged in a composite image. mCherry intensities were unchanged while brightness and contrast of the YFP channel were optimised on each plant to provide the best morphological outlines of the tissues.

### Macroscopic plant phenotyping

Macroscopic pictures were taken using different devices, according to the manufacturer’s instructions: Keyence VHX-900F, Canon EOS 450D camera, camera device of a Samsung Galaxy A10 and a Ulefone Armor X5 pro.

### Phenotypic measurements

Basal bract score was determined by counting all bracts in the main stem and cauline branches, normalising by the total number of branches (excluding rosette branches, **Fig. 1E**); the inspection was limited to the first five flowers, especially for mutants. The number of bracts was counted after internodes elongation on the last upper cauline branch to ensure bracts were visible to the naked eye. Flowering time was measured with different methods (mentioned in the main text): the number of days from the start of transfer to in growth chambers to the day of bolting, the day when the first flower blooms (open petals), or as the cumulated number of leaves (rosette only or rosette and cauline). For plastochron measurements, several plants of both genotypes were synchronised and grown in the same condition. Each sampling day after transfer to a long day, 5 to 10 plants were randomly dissected under a binocular dissecting scope, removing and counting all organs (leaves or flowers) from the first leaf to the youngest organ visible on the main meristem. The youngest organs were counted as soon as they were separated by a boundary (corresponding to stage-2 flowers as defined by (Smyth et al. 1990).

### DNA extraction and sequencing for Bulk Segregant Analysis (BSA)

For the BSA, an F_2_ mapping population of 684 plants was generated by crossing (*Col-0* x *Tsu-0*) in both directions. The bulk segregant analysis was split into four replicates. A 1cm² leaf sample was sampled for each F_2_ plant, kept at −20°C and genomic DNA was extracted individually if the plant was selected in one of the bulk. Genomic DNA of 56 and 17 plants were selected and pooled in the bulks of low and high bract-score plants, respectively. The genome of the parental lines was re-sequenced using genomic DNA from bulk *Tsu-0* and *Col-0* seedlings, respectively. All genomic DNAs were extracted and purified using a CTAB-based protocol (cetyl trimethylammonium bromide), following instructions as in (Healey et al. 2014).

Each DNA bulk was prepared by pooling the genomic DNA of selected plants in equal quantities, to reach a final concentration between 13 and 25 ng/µl. Pooled DNA bulks and parental DNA were used to prepare libraries and sequenced on BGISEQ-500WGS according to the manufacturer’s instructions, yielding 5 Gb data of 100bp paired-end reads per library (target coverage of 40X).

### DNA sequencing analysis and genomic variant analysis

Sequencing data consists of two parental plus two bulks of sequencing data. A genomic variant analysis was performed on each dataset following the workflow of short variant discovery previously described in (Besnard et al. 2020). This resulted in four gVCF files (one per sample) generated by the HaplotypeCaller tool of GATK (v3.8, McKenna et al., 2010). TAIR10 was used as the *A. thaliana* reference genome. Then, the two parental gVCF were first joint-genotyped using GATK’s tool GenotypeGVCFs to emit a common vcf file for the two samples. This file was used to select a list of specific SNPs and small indels of *Tsu-0* (91AV stock) versus *Col-0* (186AV stock), filtering for positions with coverage metric DP>10. This reference list of *Tsu-0* polymorphisms was then used as the --dbsnp option in a second pass of joint genotyping using all four gVCF as input (two parental samples plus the two bulks) to emit a common vcf file. Finally, relevant polymorphic positions from the reference list in the two bulks were selected after filtering for a depth ≥3.

### QTL mapping from Bulk Segregant Analysis (BSA)

QTL identification was carried out using the QTLseqr package (Mansfeld and Grumet 2018) according to user instructions. After filtering data with the following parameters (refAlleleFreq = 0, minTotalDepth= 10, maxTotalDepth = 90, minSampleDepth = 10, minGQ= 99, depthDifference = 15), the deltaSNP index (Takagi et al. 2013) was generated and loci that reached the approximately 95% confidence interval were retained.

### QTL mapping using Recombinant Inbred Lines

Genotyped RILs from *Col-0 x Tsu-0* (3RV) are publicly available in the VASC. Based on BSA results, a panel of 55 RIL were selected from their known genotypes on chromosomes 1 and 5 using GGT 2.0 software (van Berloo 2008). The detailed genotypes of each line used in this study are available in **Table S9**. The presence of basal bracts was quantified in each line using the bract score. QTL mapping was performed with R/qtl software according to user instructions (Broman et al. 2019): we used the scanone function with the mean value of the bract score for the 55 tested RILs as a trait and significant thresholds were computed by setting a permutation number to 2000. For the bract score, the “non-parametric” (np) statistical model was used while for paraclade number, we used a normal model implemented with the hk method. To look for QTL interactions, the scantwo function was used with the normal model and the hk method since the np model is not implemented for this function. MQM was performed with default parameters. The (broad-sense) heritability for the bract score was defined as *H^2^ = (var_T_ − var_E_) / var_T_*, with *var_T_* and *var_E_* being the total and environmental variance, respectively. *var_T_* was computed as the total variance of the bract score of each plant over the entire QTL mapping dataset (1830 RIL plants and 780 control parents split over 12 experiments) while *var_E_* was computed as the pooled variance of the line variances (computed over all plants of a line across all experiments), which averages the different line variances as is computed as: (∑_i=1,N_((n_i_ - 1) x s_i_^2^))/(∑_i=1,N_ (n_i_ - N)) where N is the number of line; n_i_ and s_i_ the number of plants in the line i. Confidence intervals of *H^2^* were provided by bootstrapping 70% of the strain 1000 times as in (Chuffart et al. 2016).

### KASP genotyping

We selected 19 new genotyping markers from the 140SNPvCol marker set (Lutz U. et al. 2017) to cover the two large QTLs mapped in chr1, 1a and 1b (**Fig. S6B**) and corresponding kompetitive allele-specific PCR (KASP) oligos were ordered to Biosearch Technologies^TM^ LGC Ltd. To further reduce the intervals, we designed two new SNP markers targeting *Tsu-0* polymorphisms identified in our whole-genome re-sequencing data. Specific KASP oligos (three per SNP) were designed by LGC from the 100-bp sequence surrounding the SNP. All 21 new markers (**Table S10**) were validated on parental genomic DNA (*Col-0* and *Tsu-0*) in a KASP assay before their use with recombinant inbred lines. Clean genomic DNA from CTAB-based extraction was used for all samples. For each marker, 2.5 µL of DNA (diluted to a final concentration of 5 ng/µL), 2.5 µL of KASP-TF V4.0 2X Master Mix low ROX and 0.07 µL KASP assay mix were mixed in 384-well plates and the genotyping assays were run in a QuantStudio 6 Flex (Applied Biosystems), using standard user guidelines for thermal cycling and final fluorescence analysis.

### Fine mapping using Heterologous inbred families

Using F_6_ genotyping data for the 3RV RIL set from the VASC, we selected 3 RILs heterozygous in a region overlapping QTL1a, QTL1b and the interval in-between: RIL 334, 488 and 478, respectively (**Fig. 4** and **Fig. S6**). For RIL 334 and 488, we genotyped by KASP 160 segregating F8 progenies and selected 5 and 8 plants, respectively, with homozygous recombination inside the QTL region under study. For RIL 478, we genotyped 20 plants and selected only one; genotyping reactions were performed using PCR primers designed to amplify the published marker of the VASC (Simon et al. 2008)(**Table S11**) and the results were read by Sanger sequencing. All selected F_8_ plants were selfed to generate an inbred line of a fixed genotyped (F_9_), the collection of F_9_ inbred lines forming a Heterologous Inbred Family (HIF). HIF lines (with at least 20 plants per line) were phenotyped for the bract score with parental control in the same experiment.

### Biological sample preparation for the transcriptome time-series profiling

After 20 days in SD, *Tsu-0* and *Col-0* meristems were dissected every day in LD conditions, to capture the precise developmental stages (especially stages T and F, see below). Three independent biological experiments were performed with 5 to 11 meristems per replicate. Stages are defined as follows: V, the day when plants were transferred to LD from SD; L, after 4 to 5 days of LD (identical for both *Col-0* and *Tsu-0,* which have the same meristem shape at this time, see **Fig. 4A**); T, the main meristem enlarges and domes, (Kwiatkowska 2008; Kinoshita et al. 2020), and lateral meristems starts being visible at the axils of young leaf primordia; F, all young organs in the meristems were identified as flowers (note that this stage occurred several days before bolting). Stage F in mutants was defined when several rounded primordia become visible at the SAM (that will become branch-like flowers or only branches). Dissections were performed from 9:00 a.m. to noon, by alternating between *Tsu-0* and *Col-0*. Micro-dissected meristems were snap frozen in liquid nitrogen and stored at −80 °C until RNA extraction. To avoid the induction of stress-related gene expression, meristem dissection did not exceed 3 min between the first organ removal and freezing. For each replicate, pooled meristems were ground with a RockyII tissue lyser according to the manufacturer’s instructions. RNA was extracted using the PicoPure RNA Isolation Kit (Arcturus, Catalog KIT0202) according to the standard protocol. RNA concentration ranged from 3 to 64 ng/µl (average 23 ng/µl), with a RIN value between 5.6 and 7.6 (average 6.8).

### RNA-sequencing

Library preparation was made using NEBNext® Ultra™ II Directional RNA Library Prep Kit for Illumina (New England Biolabs); NEBNext® Poly(A) mRNA Magnetic Isolation Module (New England Biolabs); and NEBNext® Multiplex Oligos for Illumina (Index Primers Sets 1, 2 et 3), from 40ng of RNA and sequenced using the NextSeq500 (Illumina). Sequence quality was controlled using the Sequencing Analysis Viewer, sequences that did not pass the quality filter were removed. Following QC an average of 43 million sequences per sample was achieved.

### RNA-seq analysis (pre-processing procedures)

Quality filtering using Trimmomatic (Bolger et al. 2014) retained for all samples 96% of the reads, which were aligned to the TAIR10 genome using STAR (Dobin et al. 2013) with the following parameters: non-canonical splice junctions were removed, multi-mapping reads were limited to 10, and only reads mapped once were considered to determine splicing junctions. Between 88 and 97% of the reads were mapped to a simple locus. Normalisation of read counts was performed using the R Bioconductor packages DESeq (Love et al. 2014), with the following parameters: genes for which the total number of reads was below 10 were discarded, and data were transformed with Variant Stabilising Transformation (VST) function (Anders and Huber 2010), to harmonise the variance. Consistency between biological replicates was verified using a Principal Component Analysis on all samples with VST-transformed data.

### RNA-seq analysis: Differential Gene Expression (DGE) analysis

Differential analysis was performed using the R Bioconductor package edgeR (McCarthy et al. 2012). Reads were first normalised using TMM (Trimmed mean of M-values) to reduce library-specific biases. Normalisation factors were between 0.94 and 1.049. A generalised linear model was applied for the analysis of differentially expressed genes (DEG). Three types of DEG analysis were considered: DEG at each stage between the different genotypes (**table S3**), DEG between the stage within the same genotype, and DEG across all conditions (stages and genotypes). Multiple DEG analyses were corrected using Benjamin-Hochberg correction, and genes with a p-value < 0.05 were retained.

### RNA-seq analysis: clustering gene expressions correlating with bract presence

To identify bract signature genes (see **Figure 5**), we first calculated the minimum and maximum expression levels among stages with and without bracts. If the minimum expression level in the stages with bracts was greater than or equal to the maximum expression level in the stages without bracts, the genes were classified as bract signature genes whose expression levels positively correlated with the presence of bracts. If the maximum expression level in the stages with bracts was lower than or equal to the minimum expression level in the stages without bracts, these genes were classified as bract signature genes whose expression levels anti-anti-correlated with the presence of bracts. If neither of these conditions were satisfied, genes were discarded.

### Temporal registration of gene expression dynamics

To align gene expression profiles of *A. thaliana Tsu-0* (query data) with *Col-0* (reference data), we utilised the curve registration method in the R package greatR (Kristianingsih, 2024). This approach involved shifting the gene expression profiles of *Tsu-0* across developmental stages (V, L, T, F) using shift factors ranging from −1 to 1; a stretch factor was not applied due to the two datasets being over similar times. The optimal shift parameter for each gene pair was identified by maximising the log-likelihood. The fit with the best registration factors for each gene was then compared to the fit with a non-registered model (without transformation) using the Bayesian Information Criterion (BIC) statistic. A lower BIC score for the registered model (with transformation) indicated that the gene expression dynamics of *Tsu-0* and *Col-0* were similar, and the expression profile differences could be resolved through registration. Conversely, a higher BIC score suggested that the profiles were best described by two individual curves, i.e. not aligned. Prior to transformation, gene expression levels in both datasets were centred and scaled using the z-score scaling method. The standard deviation for the replicates at each time point was set to 0.01.

### GO term analysis

GO term enrichment analyses were performed using either the ‘Single Enrichment Analysis’ tool from AgriGOv2 (Tian et al. 2017) with default parameters (Fisher’s test with Yekutieli adjustment method) or clusterProfiler 4.0 (Wu et al. 2021) with default parameters (enrichGO and simplify functions with “BH” adjustment method). The Rich factor was computed by dividing the number of genes associated with a GOterm (“Count”) by the numerator of the “BgRatio” computed by the enrichoGO function.

### Gene selection inside genetic mapping intervals

The R library ‘GenomicFeatures’ (Lawrence et al. 2013) was used to import gene information from the most recent annotation file of *A. thaliana* at the gff format (Cheng et al. 2017). Custom R scripts were used to intersect all gene loci falling within genetic mapping intervals and provide for each gene information from RNA-seq analysis and genomic variant analysis (see **Table S2** and **Table S4**). We used snpEff v5.0d (Cingolani et al. 2012) and its putative functional categories (HIGH/MODERATE/LOW/MODIFIER) to predict the functional impacts of the genomic variants over regions covering each annotated gene of the interval, including promoter (2 kbp upstream of the transcriptional start site) as well as 500-bp downstream region in the case of miRNA. In **Table S2** and **S4**, additional gene information was retrieved from ThaleMine (Pasha et al. 2020) and GO terms from the TAIR bulk data retrieval tool.

### Data handling, data visualization and descriptive statistics

The R software was used (R Core Team 2018), especially the following packages: Bioconductor packages (Huber et al. 2015) for the analysis of omics data, especially GenomicFeatures, IRange (Lawrence et al. 2013), rtracklayer (Lawrence et al. 2009), clusterProfiler (Wu et al. 2021), limma (Ritchie et al. 2015), DESseq2 (Love et al. 2014), edgeR (Chen et al. 2016) and org.At.tair.db (Carlson 2024); dplyr (Wickham et al. 2023), tidyr (Wickham et al. 2024) and reshape2 (Wickham 2007) for data manipulation and ggplot2 for almost all plots (Wickham 2016), with viridis and plasma (Garnier et al. 2024) for some colour optimization.

### Text and English

Artificial intelligence-based language correction software was used throughout the text to ensure accurate spelling and grammar, as well as to achieve the most comprehensible style possible, without influencing the content.

## Supporting information

all supplemental figures from S1 to S14

## Acknowledgements

We thank Teva Vernoux and Yoan Coudert for critical reading of the manuscript. We thank Mathieu Hanemian, Gaël Yvert and Bjorn Pieper for their advice in quantitative genetics. We thank Thomas Widiez and Nathanaël Jacquier for their advice in Kasp genotyping. We thank the following researchers for providing *A. thaliana* mutant seed stocks: Mitsuhiro Aida for *bop1-4* x *bop2-11*, and *puchi-1 x bop1-4 x bop2-11*; Hicham Chathane for *lfy-12*; Georges Coupland and Enric Bertran Garcia de Olalla for *soc1-2*. We acknowledge the PSMN (Pôle Scientifique de Modélisation Numérique) computing centre of ‘ENS de Lyon’ for providing support in the genomic variant analysis. We thank INRAE for a grant from the BAP department (call IB2019), the Embassy of France in the United Kingdom for a short-term travel fellowship awarded to R. K. R.M. and R.W. gratefully acknowledge support from the Biotechnology and Biological Science Research Council Institute Strategic Programme ‘Building Resilience in Crops’ (BB/X01102X/1). SD was supported by a Ph.D contract of the french ministry of Research and Higher Education.

## Author contributions

Sana Dieudonné: Conceptualization, Investigation, Software (implementation), Formal analysis, Visualization, Methodology, Writing -review and editing; Ruth Kristianingsih: Formal Analysis, Methodology, Software (designing, programming, testing, implementing), Visualization, Writing -review and editing; Stephanie Laine, Investigation; Béline Jesson: Formal analysis, Véronique Vidal: Supervision, Rachel Wells: Supervision, Writing -review and editing; Richard Morris: Supervision, Writing -review and editing; Fabrice Besnard: Conceptualization, Project administration, Supervision, Investigation, Formal analysis, Methodology, Software (programming, implementing), Funding acquisition, Visualization, Writing - original draft, review and editing.

## Data availability

Whole-genome DNA-seq data of the parental lines (*Col-0* and *Tsu-0*) and of the two F2 pools used for bulk segregant analysis are deposited under this identifier: doi:10.57745/Z80SIM. Time-course RNA-seq data of *Col-0* and *Tsu-0* micro-dissected meristems during flowering are available with these doi: doi:10.57745/DKMQ06, doi:10.57745/7JI3E7, respectively.

## Supporting Information

**Figures S1 to S13**

**Figure S1**: Example of natural transient bract formation at the base of branches among *Brassicaceæ* tribes.

**Figure S2**: The presence of basal bracts in *A. thaliana* accessions does not correlate with their geographic or genetic origin.

**Figure S3**: *Tsu-0* plants flower later than *Col-0* plants in different culture conditions.

**Figure S4**: Bulk segregant analysis of basal bract formation (*Tsu-0* x *Col-0*) identifies four putative QTLs.

**Figure S5**: Quantitative genetics of basal bracts formation in *Tsu-0* using a set of RILs with the reference accession *Col-0*.

**Figure S6**: Fine mapping of the QTLs controlling basal bract formation in *Tsu-0*.

**Figure S7**: QTL1b* identified for basal bract correlates with a higher cauline branch number.

**Figure S8**: Transgressive phenotypes in HIF reveal epistasis controlling floral determinacy between alleles of *Col-0* and *Tsu-0*.

**Figure S9**: Gene expression dynamics during the floral transition in micro-dissected meristems of *Col-0* and *Tsu-0* plants.

**Figure S10**: Phenotypic differences between the bracts of the mutants affected in the FMI pathway and the natural basal bracts observed in *Tsu-0*.

**Figure S11**: Functional analysis of the genes differentially expressed in *Tsu-0* meristem at the stage T when bract repression is delayed.

**Figure S13**: Evolution of the proportions of DEGs between *Tsu-0* and *Col-0* depending on the heterochronic shift and on the developmental stage

**Figure S13**: Temporal registration of expression dynamics in *Tsu-0* over the floral transition for genes related to bract development, floral identity and floral transition.

**Figure S14**: A working model for natural basal bract formation in *Arabidopsis thaliana*.

**Table S1 to S11**

**Table S1:** Literature presenting mutants in genes functionally associated with the FMI pathway in and showing derepressed bract formation.

Note that tables S2 to S11 are large spreadsheets available for download:

**Table S2**: List of annotated genes lying in the four mapped QTLs controlling bracts in *Tsu-0*, with additional information from RNAseq and genomic variant analysis.

**Table S3**: Differential gene expression analysis and log2CPM for the genes expressed in microdissected meristems of *Tsu-0* and *Col-0* during the floral transition.

**Table S4**: Intersection of the genetic mapping, the genomic variant analysis and the RNAseq data.

**Table S5:** Bract signature genes in *Tsu-0* identified by clustering apart expression from bract and non-bract making stages in *Tsu-0* and *Col-0* time-series.

**Table S6**: Details for all genes associated with “response to salicylic acid” about their expression at stage in *Tsu-0* and their identification as bract signature genes.

**Table S7**: Details for all genes associated with anthocyanin metabolism about their expression at stage in *Tsu-0* and their identification as bract signature genes.

**Table S8**: Results of curve registration

**Table S9**: Genotypes of all RIL and HIF lines used in this study.

**Table S10**: SNP information relative to new KASP genotyping markers used in this study.

**Table S11**: SNP information relative to new sanger genotyping markers used in this study and associated primers.

