## Supplementary material for "*Arabidopsis* natural variation induces complex transcriptomic heterochronies at the floral transition and perturbs the coupling between leaf and flower development by an unexpected genetic control": all supplemental figures from S1 to S14

The following Supporting Information is available for this article:

**Figures S1 to S13**

**Figure S1**: Example of natural transient bract formation at the base of branches among *Brassicaceæ* tribes.

**Figure S2**: The presence of basal bracts in *A. thaliana* accessions does not correlate with their geographic or genetic origin.

**Figure S3**: *Tsu-0* plants flower later than *Col-0* plants in different culture conditions.

**Figure S4**: Bulk segregant analysis of basal bract formation (*Tsu-0* x *Col-0*) identifies four putative QTLs.

**Figure S5**: Quantitative genetics of basal bracts formation in *Tsu-0* using a set of RILs with the reference accession *Col-0*.

**Figure S6**: Fine mapping of the QTLs controlling basal bract formation in *Tsu-0*.

**Figure S7**: QTL1b* identified for basal bract correlates with a higher cauline branch number.

**Figure S8**: Transgressive phenotypes in HIF reveal epistasis controlling floral determinacy between alleles of *Col-0* and *Tsu-0*.

**Figure S9**: Gene expression dynamics during the floral transition in micro-dissected meristems of *Col-0* and *Tsu-0* plants.

**Figure S10**: Phenotypic differences between the bracts of the mutants affected in the FMI pathway and the natural basal bracts observed in *Tsu-0*.

**Figure S11**: Functional analysis of the genes differentially expressed in *Tsu-0* meristem at the stage T when bract repression is delayed.

**Figure S13**: Evolution of the proportions of DEGs between *Tsu-0* and *Col-0* depending on the heterochronic shift and on the developmental stage

**Figure S13**: Temporal registration of expression dynamics in *Tsu-0* over the floral transition for genes related to bract development, floral identity and floral transition.

**Figure S14**: A working model for natural basal bract formation in *Arabidopsis thaliana.*

**Table S1 to S11**

**Table S1:** Literature presenting mutants in genes functionally associated with the FMI pathway in  and showing derepressed bract formation.

Note that tables S2 to S11 are large spreadsheets available for download:

**Table S2**: List of annotated genes lying in the four mapped QTLs controlling bracts in *Tsu-0*, with additional information from RNAseq and genomic variant analysis.

**Table S3**: Differential gene expression analysis and log2CPM for the genes expressed in microdissected meristems of *Tsu-0* and *Col-0* during the floral transition.

**Table S4**: Intersection of the genetic mapping, the genomic variant analysis and the RNAseq data.

**Table S5:** Bract signature genes in *Tsu-0* identified by clustering apart expression from bract and non-bract making stages in *Tsu-0* and *Col-0* time-series.

**Table S6**: Details for all genes associated with “response to salicylic acid” about their expression at stage in *Tsu-0* and their identification as bract signature genes.

**Table S7**: Details for all genes associated with anthocyanin metabolism about their expression at stage in *Tsu-0* and their identification as bract signature genes.

**Table S8**: Results of curve registration

**Table S9**: Genotypes of all RIL and HIF lines used in this study.

**Table S10**: SNP information relative to new KASP genotyping markers used in this study.

**Table S11**: SNP information relative to new sanger genotyping markers used in this study and associated primers.

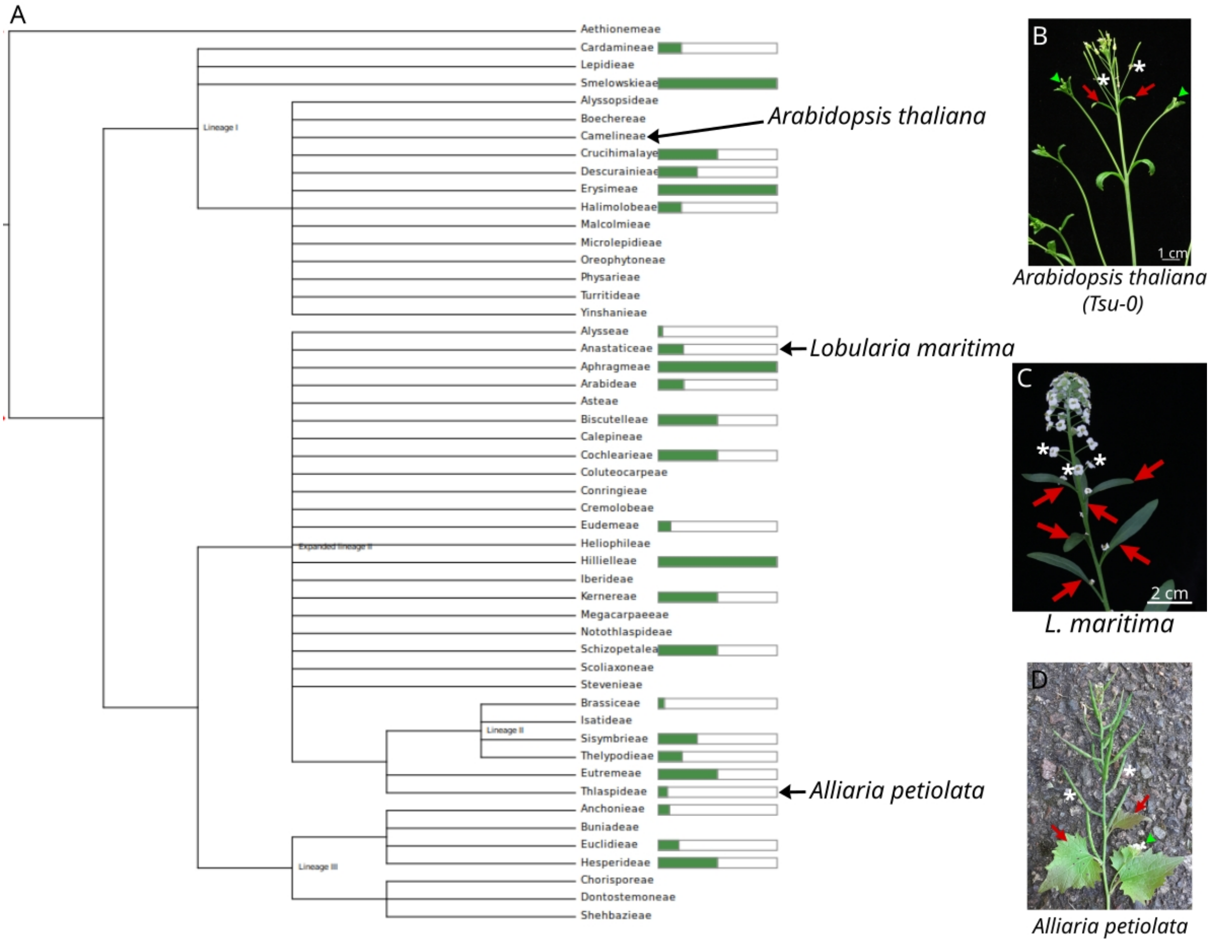


**Figure S1**: **Example of natural transient bract formation at the base of branches among *Brassicaceæ* tribes.**

**A,** Cladogram of the main *Brassicaceae tribes*: green bars at line ends indicate the number of genera per tribe with species exhibiting "*racemes that are bracteate throughout or at least in lower half*" (if absent, no such species has been reported; source: Brassibase). Such a trait can be seen in *L. maritima* (**C**) and is spread across the phylogeny with no obvious evolutionary pattern. More discrete, basal bracts in only the very first flowers (1-5) can also be observed in different species such as *A. thaliana* (**B**) or *A. petiolata* (**D**) .

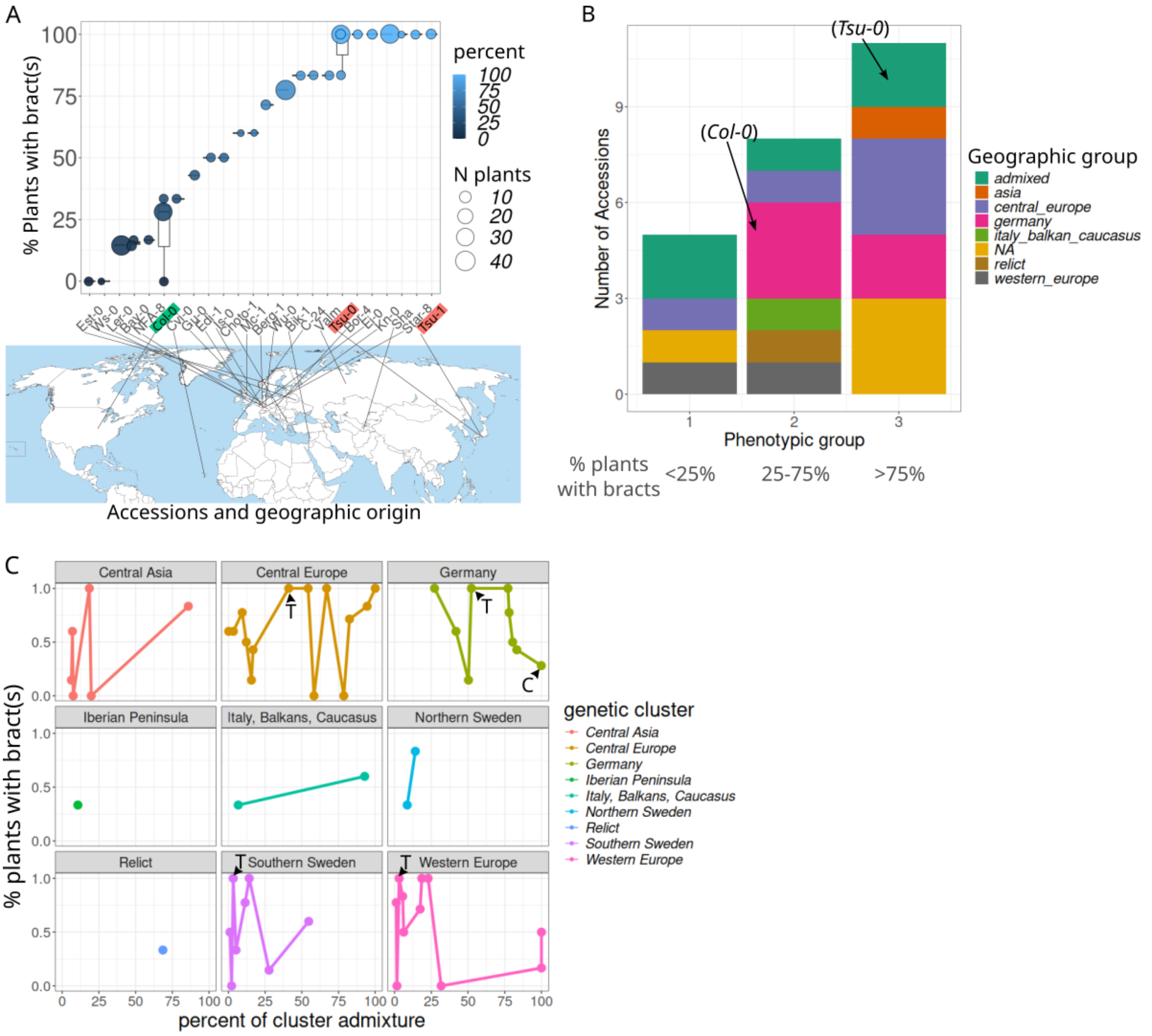


**Figure S2**: **The presence of basal bracts in *A. thaliana* accessions does not correlate with their geographic or genetic origin**

**A**, Occurrence of basal bracts in different accessions, assessed by the percentage of plants with at least one basal bract in the inflorescence. Each dot is the average value of several plants (number indicated by dot size) of a scoring assay. A box plot indicates several assays per line and thicker horizontal black lines are the median value of all scoring assays (range 1-3) in the accession. *Col-0* and *Tsu* are highlighted in green and red, respectively. The geographical origin of each strain is located in the world map below.

**B**, Accessions screened in **A** are split into three discrete groups based on bract frequency and the geographic group of the corresponding accessions is displayed as stacked bars. The positions of *Col-0* and *Tsu-0* accessions in this classification are indicated with arrows. Each phenotypic group is composed of many geographic origins.

**C**, Correlation study between the frequency of plants with bract(s) in percent (data from **A**) and the genetic composition of the accession modelled as an admixture of the 9 main worldwide genetic clusters (source: 1001 genome). T and C indicate *Tsu-0* (that is an admixture of 4 genetic clusters) and *Col-0* (that is a pure genetic cluster) for each genetic cluster, respectively. No obvious correlation appears between a genetic cluster and the presence of bracts.

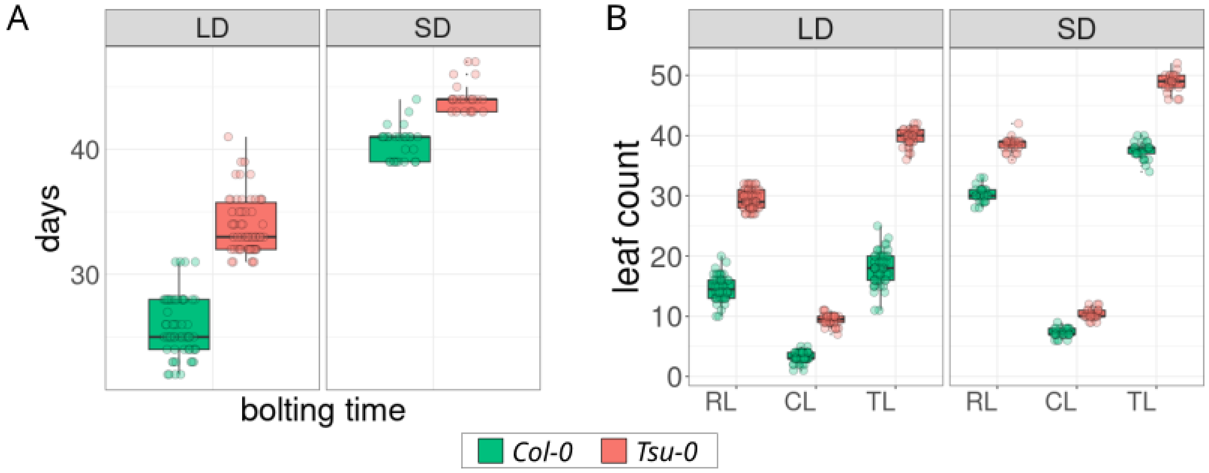


**Figure S3**: ***Tsu-0* plants flower later than *Col-0* plants in different culture conditions.** Differences in flowering time between *Col-0* and *Tsu-0* accessions are measured as bolting time in days (**A**) or as the number of leaves (**B**; RL: rosette leaves, CL: cauline leaves of branches, TL: total leaves) on the main stem. Plants grown in LD conditions (N = 58 plants per genotype split into two independent replicates) or in SD conditions (N > 23 plants, one replicate).

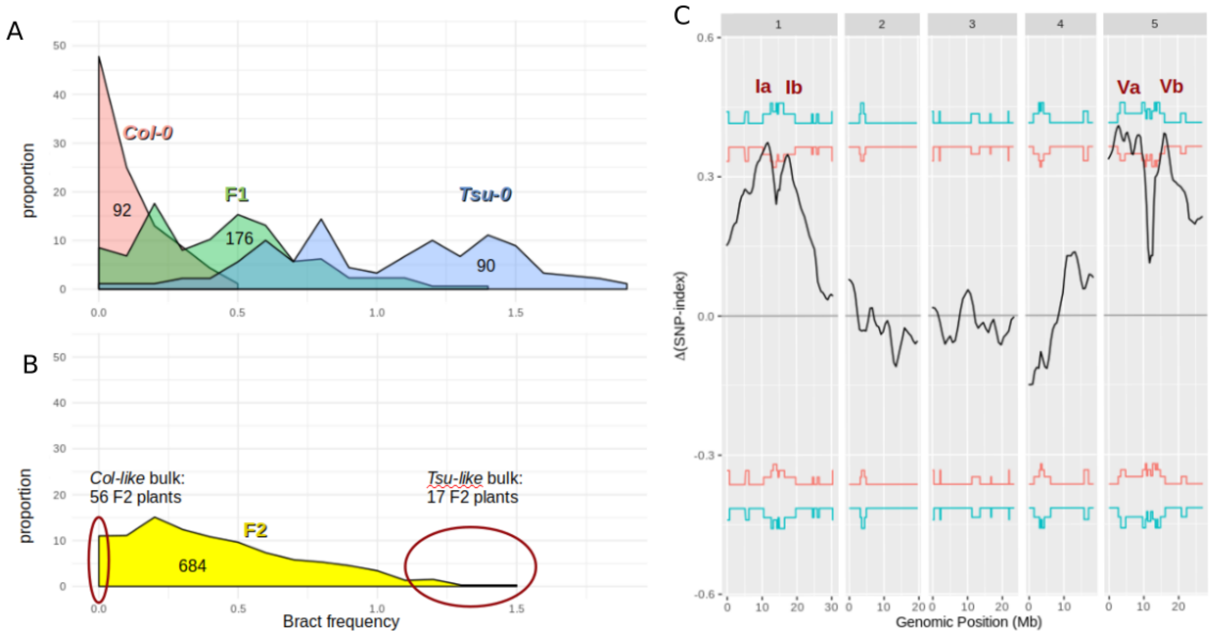


**Figure S4**: **Bulk segregant analysis of basal bract formation (*Tsu-0* x *Col-0*) identifies four putative QTLs**.
**A**, Distribution of bract scores in *Col-0* (red) and *Tsu-0* (blue) parents and in a population of F1 hybrids (green, see also figure 3A). F1 plants are a mix of crosses in both directions.
**B**, Distribution of bract scores in a population of F2 hybrids plants (see also figure 3A). The red circles indicate the plants at both tails of the distribution selected and pooled for the bulk segregant analysis (BSA).
**C**, Result of the BSA using the ΔSNP index (Takagi et al. 2013). Red and blue lines indicate the 95% and 99% confidence intervals associated with each position. The four significant peaks are labelled as Ia, Ib (on chromosome 1), Va and Vb (on chromosome 5).

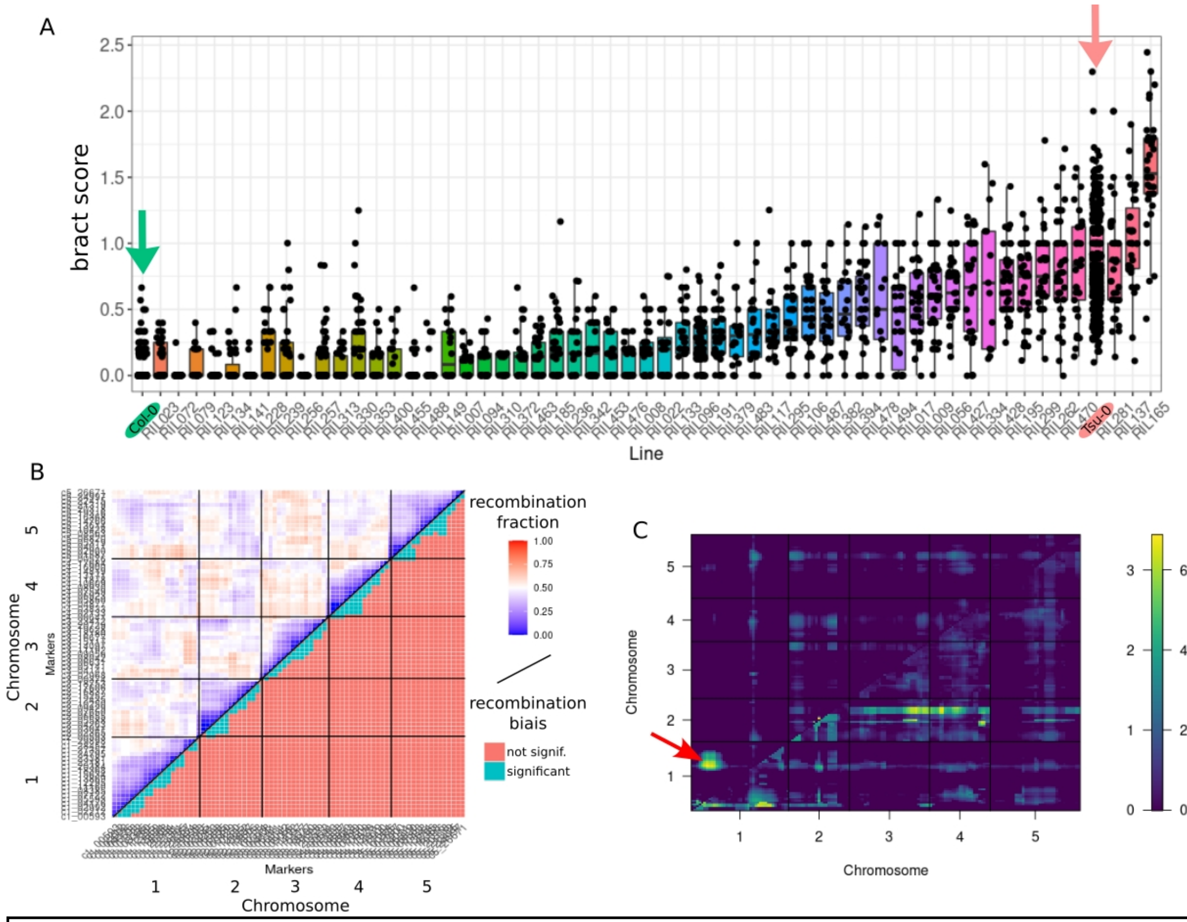


**Figure S5**: **Quantitative genetics of basal bracts formation in *Tsu-0* using a set of RILs with the reference accession *Col-0***.

**A**, Boxplots of bract scores among the 55 RILs used in this study. Parental lines *Col-0* and *Tsu-0* are highlighted in green and red, respectively and pointed with a vertical arrow of the same color. Each black dot is a plant, all lines have been scored in different assays one or several times but all assays are pooled here (1830 RIL plants in total). *Col-0* and *Tsu-0* were systematically scored in all assays as controls (780 plants in total).

**B**, Upper triangle: heatmap of the recombination fraction between the 79 genetic markers in the selected RIL set shown in A (genotyping data from Versailles stock center). Lower triangle: heatmap of the result of a Fisher test with fdr correction for multiple testing indicating whether the recombination fraction between two markers is biased on either side (too low or too much). Only the intra-chromosomal deficit of recombination is significant (fdr threshold: 0.1), as expected.

**C**, r/qtl scantwo plot. The upper triangle plots the evidence for a second additive QTL, assuming no epistasis (compared to the best single QTL-model), while the lower triangle allows for epistasis. Red arrow: there is a significant additive interaction between the two peaks of chromosome 1 (1a and 1b, p.value=0.001). Note that only the “hk” method is available for this computation, while the QTL mapping of Figure 3B is performed using the “np” model, which better fits the observed distribution of bract scores (see methods).

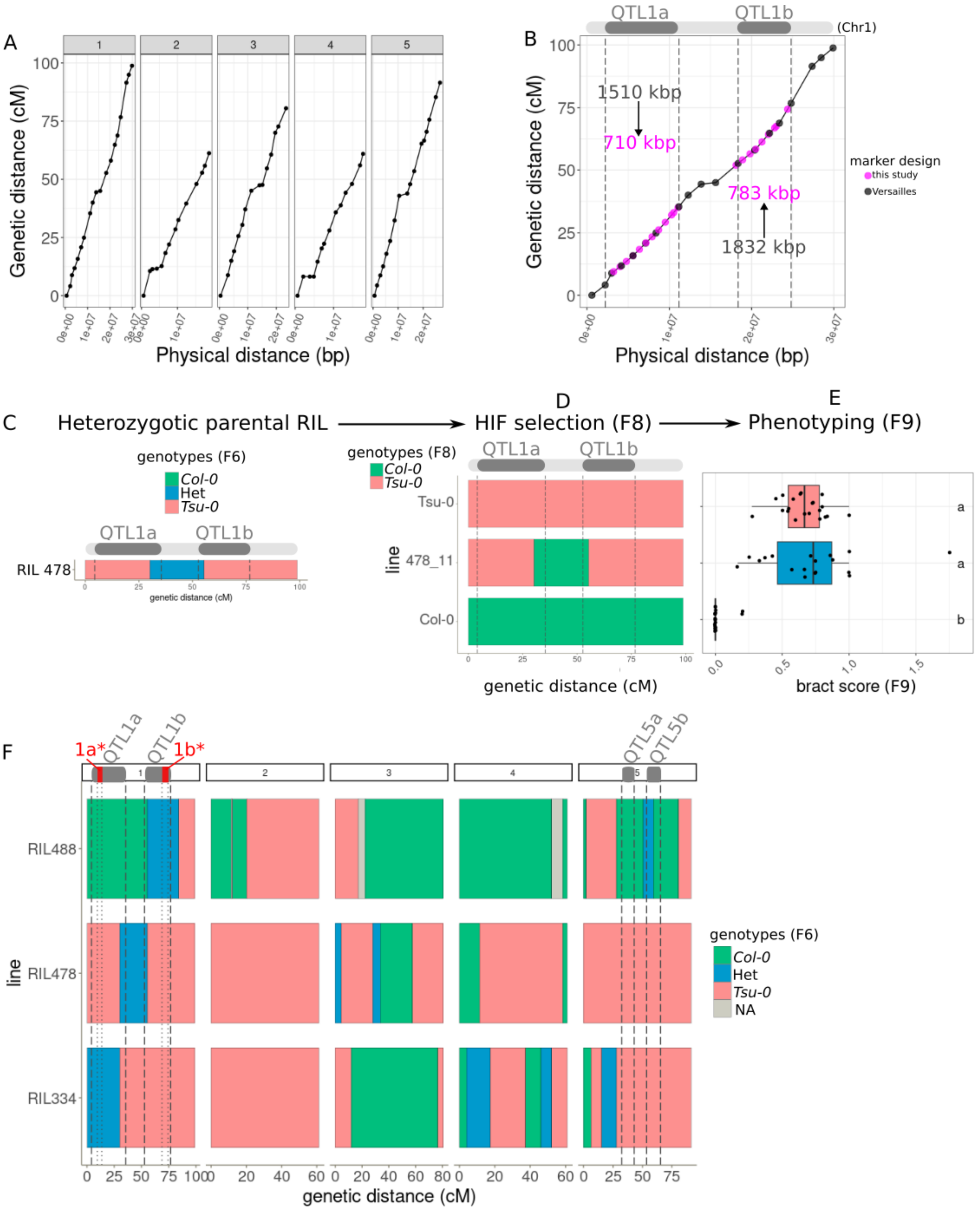


**Figure S6**: **Finer mapping of the QTLs controlling basal bract formation in *Tsu-0***.
**A**. Marey map (genetic versus physical distance) in the entire *Tsu-0* x *Col-0* RIL set (276 lines) using the 79 genotyping markers provided by Versailles resource center (Simon et al. 2008).
**B**, Marey map of chromosome 1 comparing Versailles genotyping markers (black dots) and the new set generated for this study (magenta). The positions of QTL1a/b are indicated by the bar above the plot and delineated with vertical dashed lines. The average interval length in these two regions is reported on the plot before (in black) and after (in yellow) the use of the new markers.
**C**, F6 genotype of the line RIL478 in chromosome 1, showing a heterozygotic region

between QTL1a and QTL1b.
**D**, Chromosome 1 genotype of a line (478_11) selected for being homozygous *Col-0* in the inter-QTL region and *Tsu-0* elsewhere.
**E**, Box plot of 478_11 bract score with parental controls (*Col-0*: green, *Tsu-0*: red, N > 18 plants per line). Lines sharing the same letter(s) are not statistically different (posthoc Tukey analysis with 0.05 sign. level from a glm of bract scores fitted with a quasi-poisson distribution).
**F**, Genome-wide genotype of the F6 generation for RIL 334, 488 (see figure 3) and 478 (this figure). QTL positions in chromosomes 1 and 5 are indicated by vertical dashed lines. Plotted from data provided by the Versailles resource center.

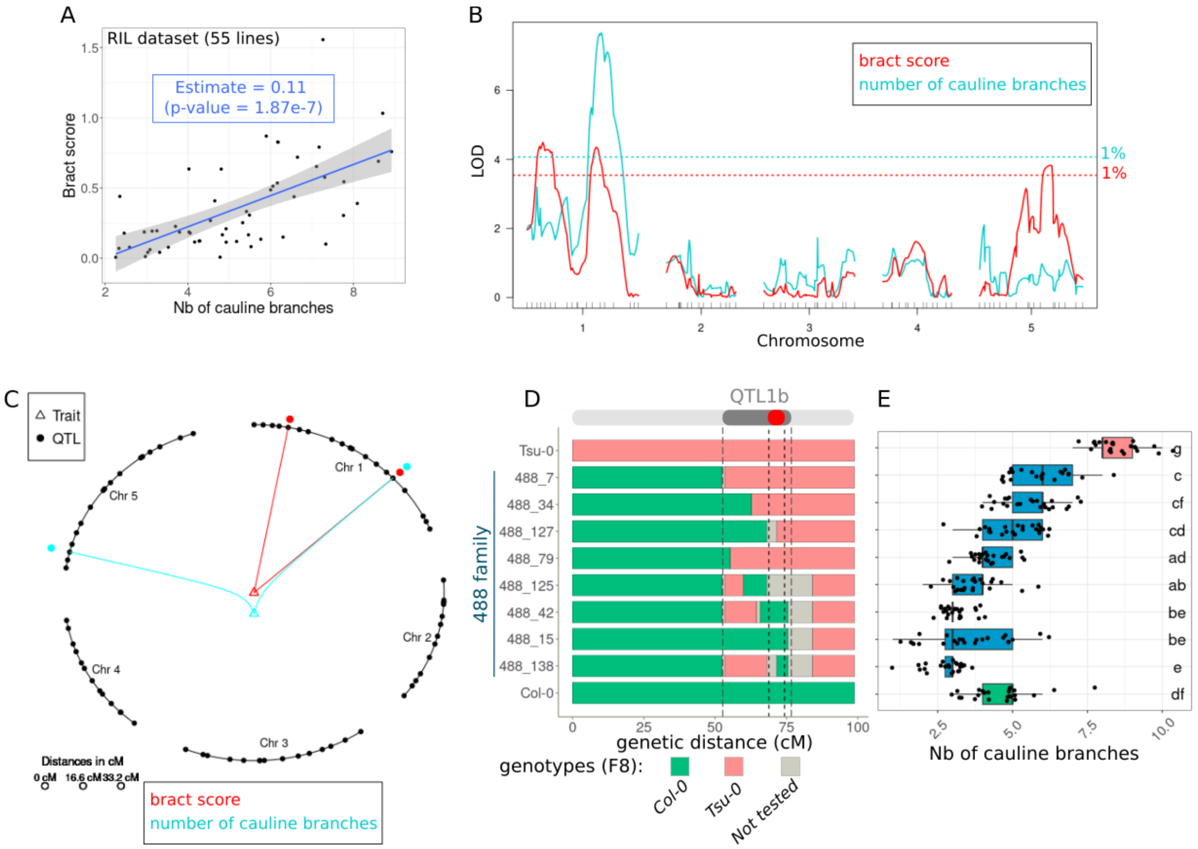


**Figure S7**: **QTL1b* identified for basal bract correlates with a higher cauline branch number**.

**A**, Plotting the average bract score against the average number of cauline branches in the 55 assayed RILs, reveals a significant correlation (blue line, standard deviation in grey). **B**, QTL mapping overlay of bract score (red) and of the average cauline branch number (blue) in the RIL set. Corresponding 1% significant thresholds after 2000 permutations are indicated by coloured dashed lines.

**C**, Using another QTL mapping method (r/qtl MQM, see method), bract score and average cauline branch number show a common QTL mapped on the same marker at the end of chromosome 1.

**D**, Chromosome 1 genotypes of selected HIF lines obtained from RIL488 (F8 generation, see figure 3). The regions QTL1b and QTL1b* mapped for bract score (see Figure 3) are delineated with vertical dashed lines.

**E**, box plots of the cauline branch number of the previous lines (blue boxes), with parental controls (*Col-0*: green, *Tsu-0*: red, N > 21 plants per line). Lines not sharing the same letter(s) are statistically different (post-hoc Tukey analysis with 0.05 sign. level from a glm of log10-transformed average cauline branch count). The *Tsu-0* allele of QTL1b* is also associated with a higher average cauline branch number.

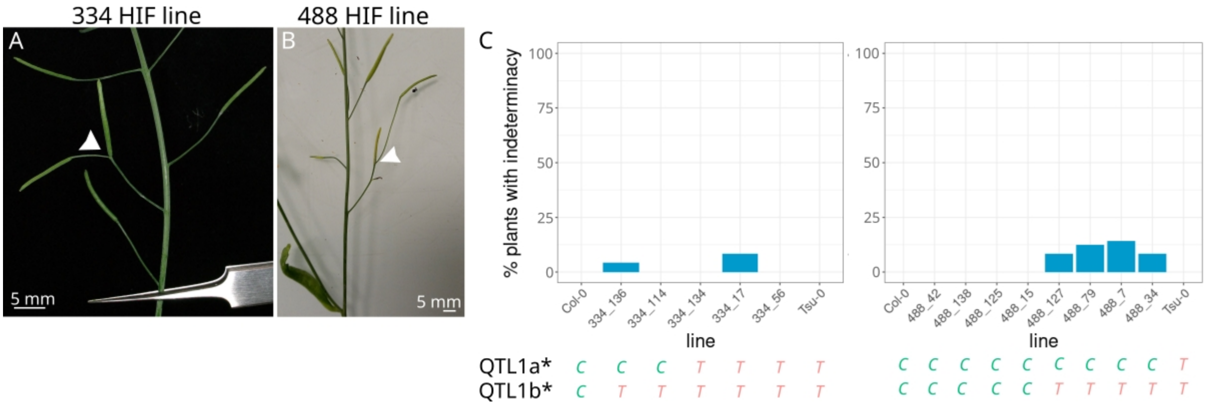


**Figure S8**: **Transgressive phenotypes in HIF reveal epistasis controlling floral determinacy between alleles of *Col-0* and *Tsu-0***.
(**A, B**) “branched” flowers (white arrowheads) observed in some RILs and HIF lines, indicating a partial loss of indeterminacy during flower development.
**C**, bar plots showing the percent of plants with indeterminacy phenotypes like in A-B among two HIF, 334 lines (left) and 488 lines (right). N > 20 plants per line. Below the bar plots, the letter indicates for each line the allelic status of QTL1a* and QTL1b* identified in Figure 3 (C, and T are *Col-0* and *Tsu-0* alleles, respectively).

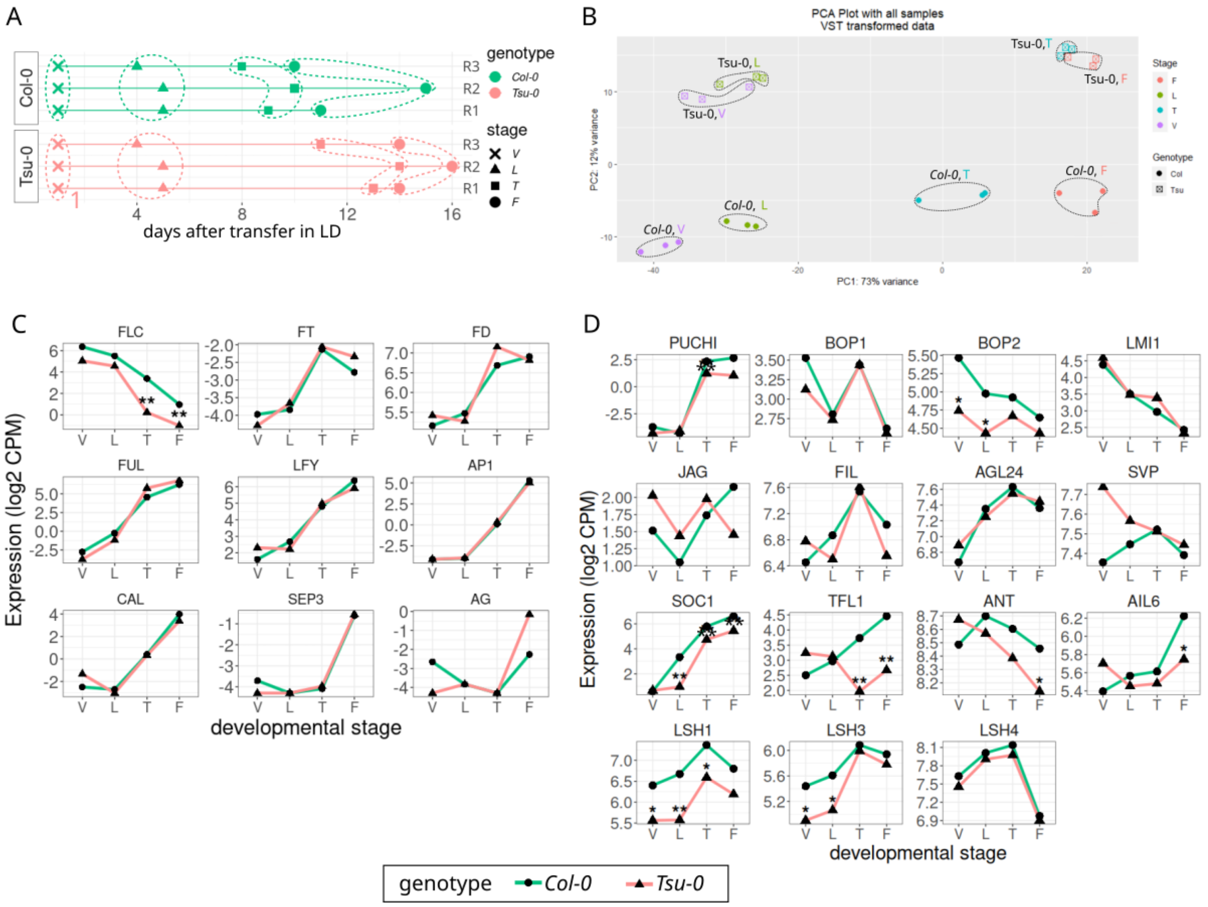


**Figure S9**: **Gene expression dynamics during the floral transition in micro-dissected meristems of *Col-0* and *Tsu-0* plants.**

**A**, Collection times (in days after transfer to long days) of the samples used in the RNAseq experiment (Fig. 5A) with corresponding developmental stages, among different biological replicates (R1-R3).

**B**, PCA on RNAseq data showing the reproducibility of the biological replicates (three per sample).

**C**, Dynamics of the expression levels of several genes controlling floral transition and identity, in *Col-0* (green) and *Tsu-0* (red).

**D**, Dynamics of the expression levels of genes reported to be involved in bract development, in *Col-0* (green) and *Tsu-0* (red).

In **C-D**, one star indicates a significant difference between the two expression levels and two stars indicate that the absolute fold change of this difference is superior to 1.

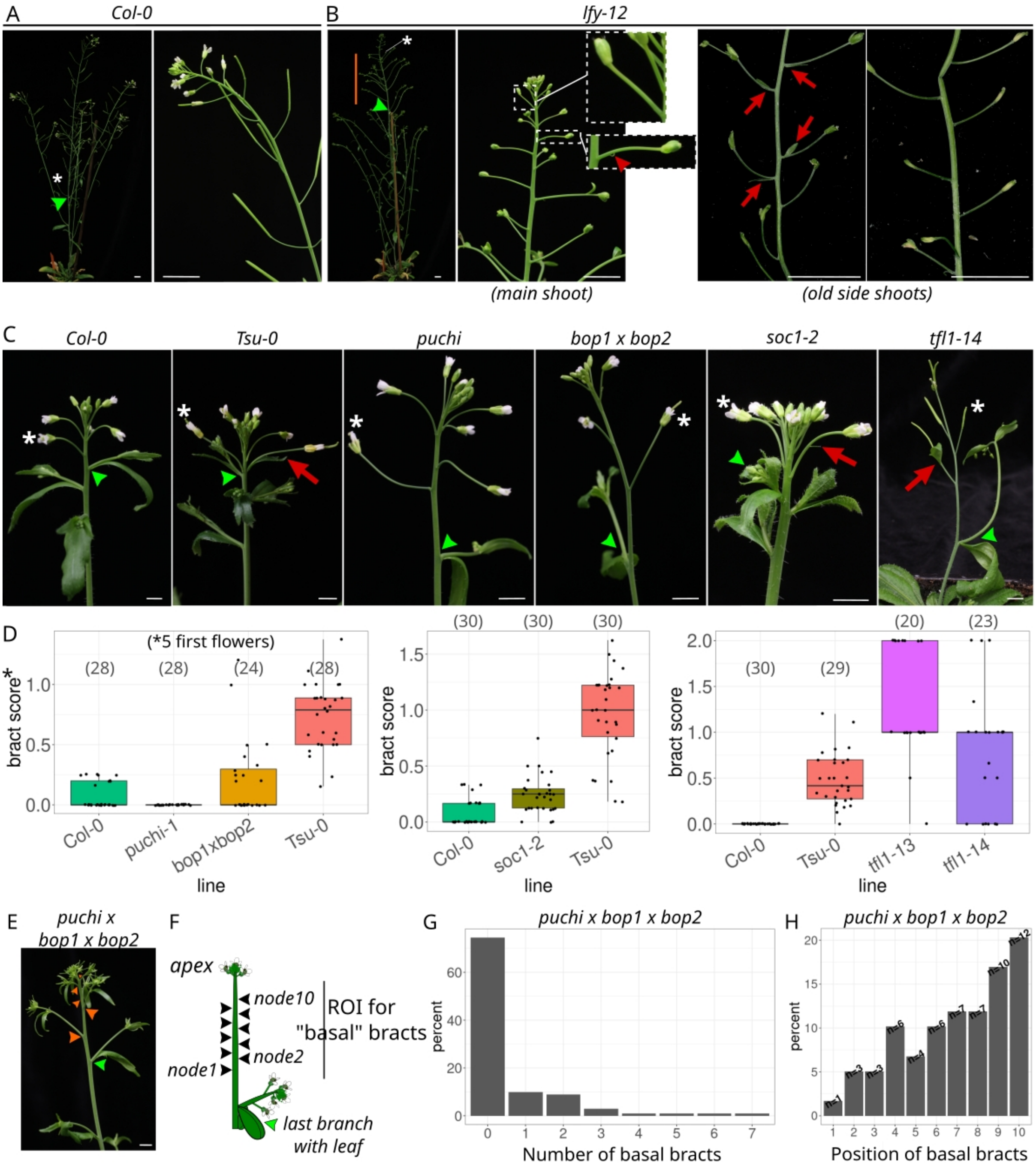


**Figure S10**: **Phenotypic differences between the bracts of the mutants affected in the FMI pathway and the natural basal bracts observed in *Tsu-0*.**
**(A, B)** Inflorescence details of a *Col-0* (**A**) versus a null *lfy* mutant plant (**B**). In wild-type *Col-0*, the last cauline branch (green arrowhead) is immediately followed by a first ebracteate flower (white star, here at mature fruit stage). In *lfy-12*, the number of normal cauline branches with leaf is increased and the last branch is followed by a region of nodes bearing I* branches (vertical orange bar), which are lateral branches with indeterminate growth and *without* leaf, appearing just after normal cauline branches but before the production of determinate floral structures (Ratcliffe et al. 1998). The first ebracteate, determinate (but mutant) flower is marked with a white star. On the main shoot, mutant flowers are mostly ebracteate, although they sometimes present rudimentary bracts (inset, red arrowhead). Fully developed bracts (red arrows) are more frequent on old axillary side shoots (right pictures), although with significant variability (note the difference between the two branches).

**C,** Pictures of the floral transition in the main stem of different lines (indicated above each picture), including wild types and classical bract mutants of the FMI pathway. The last cauline branch is indicated with a green arrowhead, the first ebracteate flower with a white star and bracts or rudimentary bracts with a red arrow.
**D**, Quantification of basal bracts using the bract score (see **Fig. 2B**) in wild-types and bract mutants for some mutants of the FMI pathway: *puchi* and *bop1 x bop2*, *tfl1-13* and *tfl1-14,* and *soc1-2*. As for *Tsu-0*, only the five first flowers are scored, but note that the *bop1 x bop2* mutant may display more bracts in later flowers. Separated panels are three independent experiments, brackets: number of plants scored.
**E**, The main stem of a *puchi x bop1 x bop2* mutant plant showing the transition between the last normal branch with a cauline leaf (green arrowhead) and several I* branches (orange arrowheads, see **A,B** legends for the definition of I* branches)
**F**, Drawing explaining how basal bracts are scored in the *puchi x bop1 x bop2* triple mutants (see panel F-G): only the ten first nodes after the last normal branch are considered (region of interest or ROI); scored shoots are the main and secondary shoots.
**G**, Distribution (as percent) of the number of bracts per branch in the ten first nodes after the last branch with a true leaf in *puchi x bop1 x bop2* plants (N plants= 21, N branches=102): Most branches (>75%) display no bract because most nodes bear I* branches (orange arrowheads in **E**).
**H,** Distribution (as percent) of the position of bracts in the ten first nodes after the last cauline branch in *puchi x bop1 x bop2* plants. When basal bracts are found in this region, their frequency increases with the distance to floral transition, the opposite trend observed for basal bracts in Tsu-0 (see **Fig. 1B**).

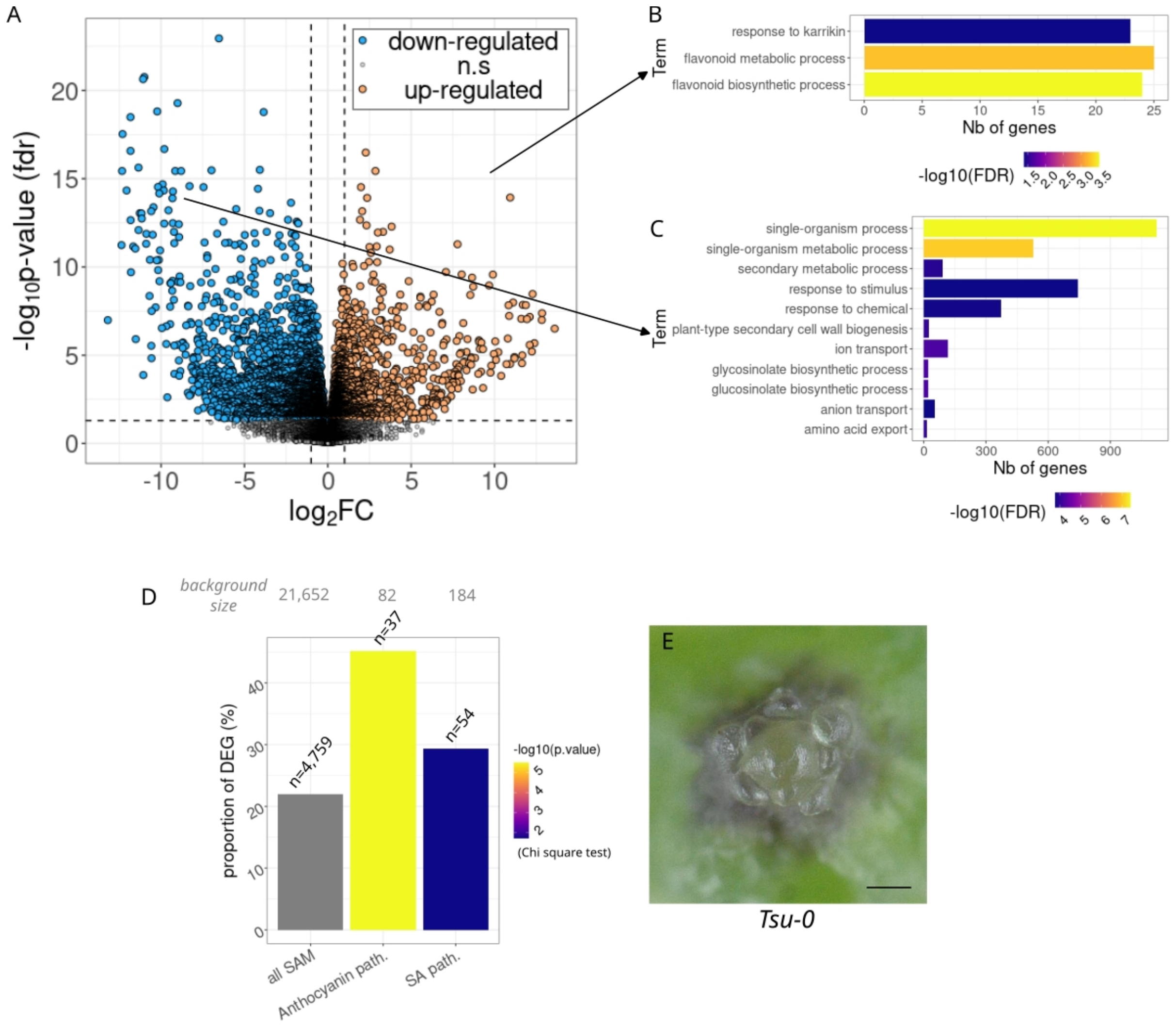


**Figure S11: Functional analysis of the genes differentially expressed in *Tsu-0* meristem at the stage T when bract repression is delayed.**

**A,** Volcano plot of gene expressions at the T stage between the two accessions. All genes expressed in the SAM are plotted (n=21,652 grey dots),genes above the statistically significant threshold (horizontal dashed line at $5.10^{-2}$) are highlighted in orange and blue for up- and down-regulation, respectively (n=4,759). Vertical dashed lines: absolute fold change superior to 1.

**B**, The only three significantly enriched 'biological process' GO terms from the DEG up-regulated in A and the corresponding number of genes.

**C**, The top-ten significantly enriched 'biological process' GO terms from the DEG down-regulated in A and the corresponding number of genes.

In B,C, color gradients indicate the statistical significance of the GO term enrichment (Fisher's test with Yekutieli adjustment method)

**D**, At the T stage, comparing the DEG ratio among all the genes expressed in the SAM versus the anthocyanin biosynthesis and SA-responding pathways indicate these groups have a significantly higher proportion of genes with a different expression between *Tsu-0* and *Col-0*.

**E**, A wild-type *Tsu-0* micro-dissected meristem at T stage showing high anthocyanin colouration at the base of the stem and up to young developing organs. scale bar=100 µm.

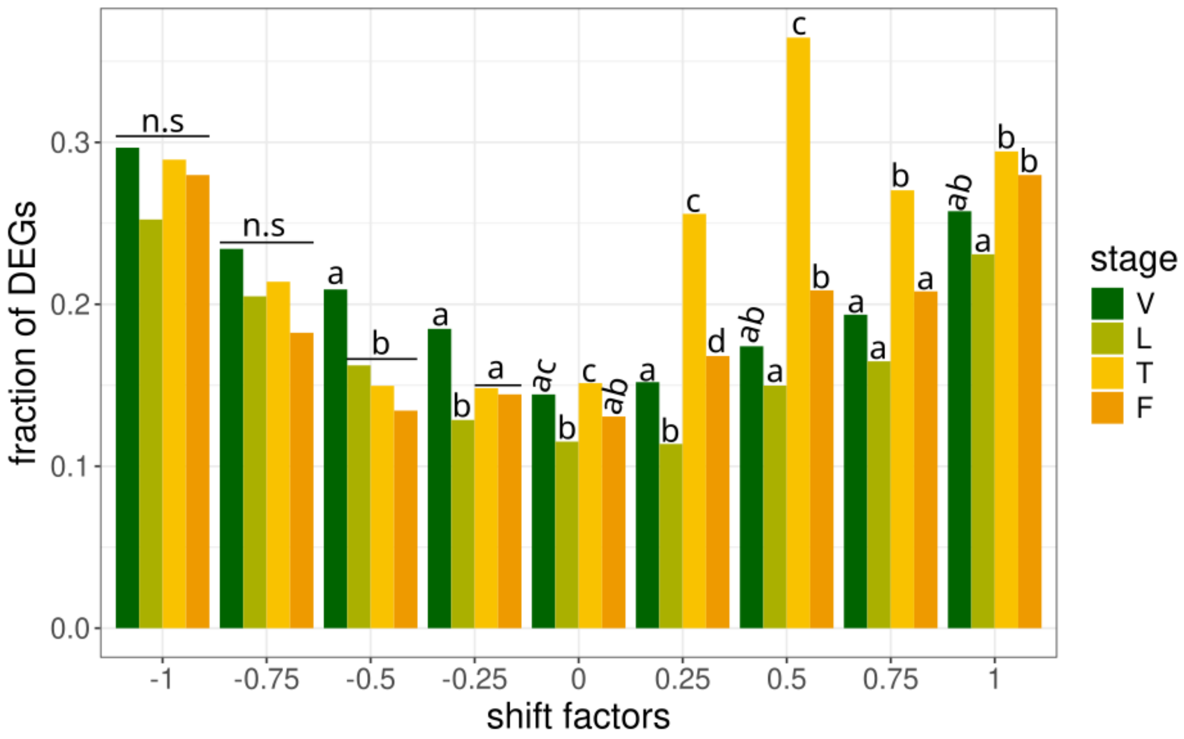


**Figure S12**: **Evolution of the proportions of DEGs between *Tsu-0* and *Col-0* depending on the heterochronic shift and on the developmental stage.**
This figure plots the same data presented in Fig.6D as a bar plot for clarity. *Post hoc* analysis of pairwise chi-square tests for a difference in the proportion of DEGs between different stages at each shift factor value (independent tests). Holm correction is applied to correct multiple testing between stages.

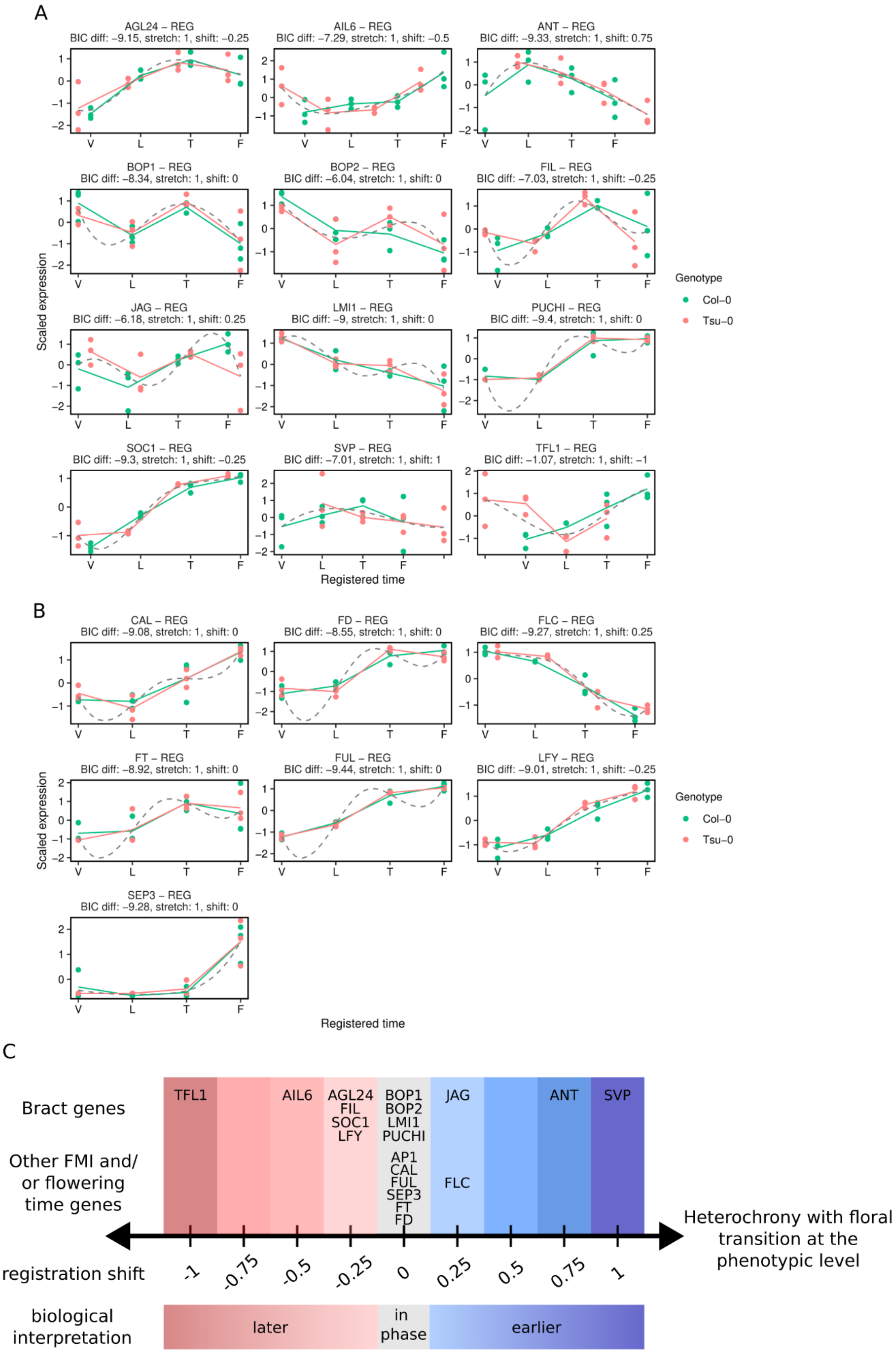


**Figure S13**: **Temporal registration of expression dynamics in *Tsu-0* over the floral transition for genes related to bract development, floral identity and floral transition.**

**A,** selection of previously known “bract” genes (see figure S12D) and the results of the temporal registration of their scaled expression dynamics between *Col-0* (green, taken as a reference) and *Tsu-0* (red).
**B**, selection of several key genes controlling floral transition and identity and the results of the temporal registration of their scaled expression dynamics between *Col-0* (green, taken as a reference) and *Tsu-0* (red).
**C**, summary of the different shift factors computed after temporal registration of genes related to bract development and/or floral transition and identity between *Col-0* and *Tsu-0* (see Figure 6 and Fig. S15 A, B)

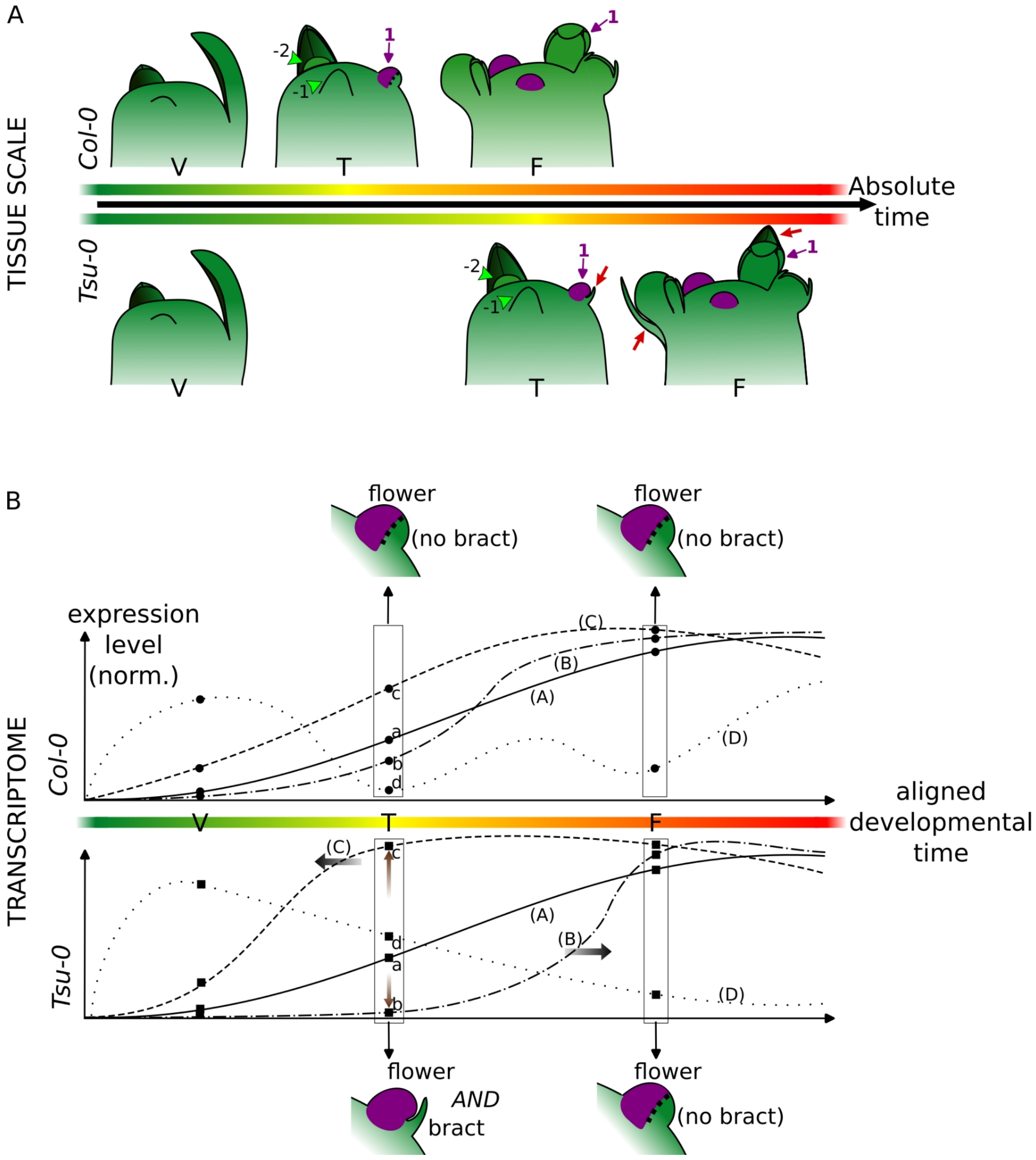


**Figure S14**: **A working model for natural basal bract formation in *Arabidopsis thaliana.***
**A**, Transition from the vegetative stage (V, producing leaves) to flowering (F, producing flowers) at the main shoot apex in *Col-0* and *Tsu-0* (top and lower row, respectively). At the transition stage (T), the first flower is formed (pointed with the purple arrow and number 1), in a sharp transition with previous axillary meristems (green arrowheads; numbered in decreasing order from the first flower). In *Tsu-0*, the first flowers co-develop with a bract (red arrows) despite the expression of floral meristem identity genes in the floral meristem (purple). In *Col-0*, the bract never outgrows from the cryptic bract domain. Rapidly, *Tsu-0* stops producing bracts. The stages happen at different absolute times (*Tsu-0* flowers later), and the developmental progression from V to F is highlighted in a tragreen-to-red color gradient for each accession.
**B**, Desynchronisation of gene expression dynamics creates a new gene expression state in

*Tsu-0*. Four hypothetical genes are depicted (their particular temporal evolution does not matter here), in *Col-0* (top) and *Tsu-0* (bottom). A is a gene synchronized with flowering transition in both accessions (e.g. *LFY* or *AP1*, see Figure 5), B's dynamics is shifted late, and C's dynamic is shifted early. Horizontal grey arrows highlight these transcriptional heterochronies for B and C genes in *Tsu-0* compared to *Col-0* and brown vertical arrows highlight the different expression levels resulting at the T stage in *Tsu-0* for those two genes (the small letters a-d label which gene's curve belongs each time point). Pure heterochronic shifts cannot explain all differences in gene expressions between the two accessions, as exemplified by the hypothetical gene D's dynamics. Altogether, the new gene expression state resulting at the T stage in *Tsu-0* promotes both flower and bract development. This state is transient and bract inhibition is restored as the meristem progresses towards the F stage.
