## Supplementary material for "*Arabidopsis* natural variation induces complex transcriptomic heterochronies at the floral transition and perturbs the coupling between leaf and flower development by an unexpected genetic control": Table S1

**Table S1: Literature presenting mutants in genes functionally associated with the FMI pathway in *Arabidopsis thaliana* and showing derepressed bract formation.**

| Mutant with derepressed bract | Corresponding gene names | references |
| --- | --- | --- |
| *lfy* | LEAFY (LFY) | Schultz and Haughn 1991; Weigel et al. 1992 |
| *tfl1* | TERMINAL FLOWER 1 (TFL1) | Shannon and Meeks-Wagner 1991; Penin 2008 |
| *ufo* | *UNUSUAL FLORAL ORGANS (UFO)* | Levin and Meyerowitz 1995; Hepworth et al. 2006 |
| *ap1* | *APETALA1* (*AP1*) | Irish and Sussex 1990; Bowman et al. 1993 |
| *bop1 x bop2*  *bop1 x bop2 x lfy*  *bop1 x bop2 x ap1*  *bop1 x bop2 x lmi1*  *bop1 x bop2 x puchi*  *bop1 x bop2 x agl24* | *BLADE ON PETIOLE1/2 (BOP1/2)*  *LATE MERISTEM IDENTITY 1 (LMI1)*  *PUCHI*  *AGAMOUS-LIKE 24 (AGL24)* | Hepworth et al. 2005; Norberg et al. 2005; Karim et al. 2009; Xu et al. 2010; Chahtane et al. 2018 |
| *soc1 x agl24 x svp* | *SUPPRESSOR OF CONSTANS 1 (SOC1)*  *SHORT VEGETATIVE PHASE (SVP)* | Liu et al. 2009 |
| *ful*  *ful x soc1* | *FRUITFUL (FUL)* | Melzer et al. 2008; Balanzà et al. 2014 |
| *fil*  *fil ufo*  *fil lfy* | *FILAMENTOUS FLOWER (FIL)* | Levin and Meyerowitz 1995; Sawa et al. 1999; Siegfried et al. 1999 |
| *lsh1 x lsh3 x lsh4* | *LIGHT-­DEPENDENT SHORT HYPOCOTYLS 1/3/4* | Rieu et al. 2024 |
